# Glutamine Deprivation Regulates the Origin and Function of Cancer Cell Exosomes

**DOI:** 10.1101/859447

**Authors:** Shih-Jung Fan, Benjamin Kroeger, Pauline P. Marie, Esther M. Bridges, John D. Mason, Kristie McCormick, Christos Zois, Helen Sheldon, Nasullah Khalid Alham, Errin Johnson, Matthew Ellis, M. Irina Stefana, Cláudia C. Mendes, S. Mark Wainwright, Christopher Cunningham, Freddie C. Hamdy, John F. Morris, Adrian L. Harris, Clive Wilson, Deborah C. I. Goberdhan

## Abstract

Exosomes are secreted extracellular vesicles (EVs) carrying diverse cargos, which can modulate recipient cell behaviour. They are thought to derive from intraluminal vesicles formed in late endosomal multivesicular bodies (MVBs). An alternate exosome formation mechanism, which is conserved from fly to human, is described here, with exosomes carrying unique cargos, including the GTPase Rab11, generated in Rab11-positive recycling endosomal MVBs. Release of these exosomes from cancer cells is increased by reducing Akt/mechanistic Target of Rapamycin (mTORC1) signalling or depleting the key metabolic substrate glutamine, which diverts membrane flux through recycling endosomes. The resulting vesicles promote tumour cell proliferation and turnover, and modulate blood vessel networks in xenograft mouse models *in vivo*. Their growth-promoting activity, which is also observed *in vitro*, is Rab11a-dependent, involves ERK-MAPK-signalling and is inhibited by antibodies against Amphiregulin, an EGFR ligand concentrated on these vesicles. Therefore, glutamine depletion or mTORC1 inhibition stimulates release of Rab11a-exosomes with pro-tumorigenic functions, which we propose promote stress-induced tumour adaptation.

## INTRODUCTION

Extracellular vesicles (EVs), produced in intracellular compartments or by plasma membrane shedding events, have emerged as critical players in cell-cell communication (Tkach and Théry, 2016, Maas et al., 2017). They deliver specific combinations of proteins, nucleic acids, and lipids to recipient cells. EVs function in normal physiological processes, such as reproduction (Corrigan et al., 2014), immune responses (Bruno et al., 2015), neural development and maintenance (Krämer-Albers and Hill, 2016), and metabolism (Thomou et al., 2017). They also have roles in pathological events (Maas et al., 2017; Huang-Doran et al., 2017; Veerman et al., 2019) with much focus on the role of EVs in promoting tumour growth, survival, and metastasis (Becker et al., 2016). These effects involve interactions of tumour cells with each other and surrounding stromal cells (Wendler et al., 2017). EVs can alter the tumour microenvironment, for example, by promoting endothelial network formation (Sheldon et al., 2010). In turn, microenvironmental stresses, such as hypoxia (Kucharzewska et al., 2013), can affect this signalling by driving changes in the tumour EV profile. The functional relevance of stress-induced EV signalling, however, has not been extensively characterised. This is of particular interest in cancer, where depletion of oxygen and key metabolites, such as glutamine (Zhang et al., 2017), is an inevitable outcome of rapid tumour growth.

Preparations of secreted EVs from cell lines and primary cell cultures include microvesicles derived from the plasma membrane and exosomes made inside cells (Théry et al., 2018). Exosomes are EVs of around 30-150 nm in diameter, formed by the inward budding of the limiting membrane of intracellular compartments, widely thought to be late endosomes. Exosome secretion results from fusion of the resulting multivesicular bodies (MVBs) with the plasma membrane (Maas et al., 2017). Endosomal markers, such as the transmembrane tetraspanins CD63 and CD81, are typically used to identify exosomes (Kowal et al., 2016; Mateescu et al., 2017). Members of the Endosomal Sorting Complexes Required for Transport (ESCRT) family (Colombo et al., 2013), and ceramides (Trajkovic et al., 2008) regulate two proposed exosome biogenesis pathways. Several endosomal Rab GTPases, which promote trafficking between specific intracellular compartments, also play important roles (Ostrowski et al., 2010). Whether these mechanisms contribute to the heterogeneity observed in exosome preparations (Zhang et al., 2018) remains largely undetermined.

We have developed a *Drosophila* model to investigate exosome biogenesis *in vivo* (Corrigan et al., 2014; reviewed in Wilson et al., 2017). This uses the prostate-like secondary cells (SCs) of the fly male accessory glands (AGs), which secrete exosomes in to the AG lumen and have unusually large intracellular membrane-bound compartments (Figure 1A), including Rab11-positive endosomes and Rab7-positive lysosomes.

**Figure 1.**
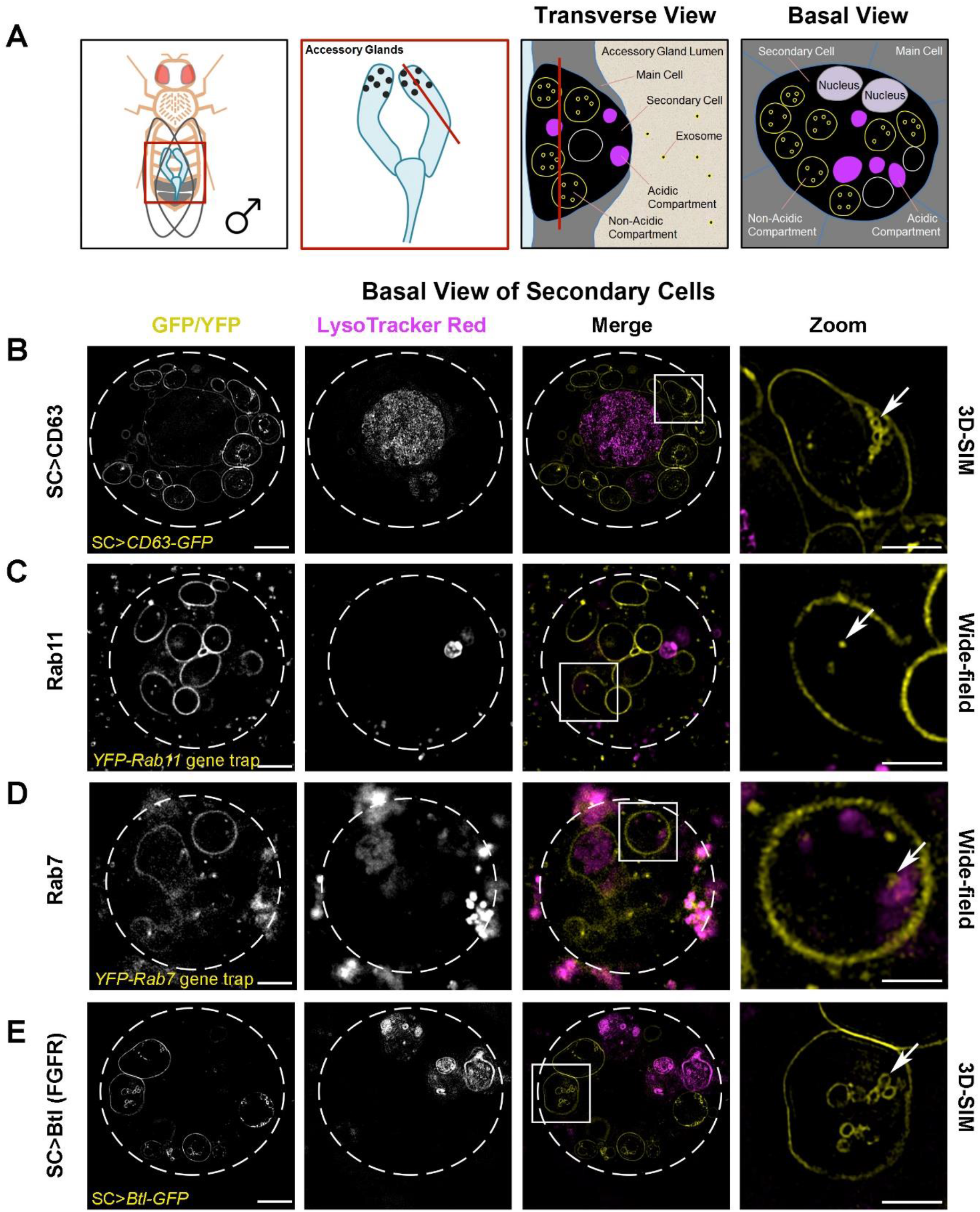
Rab11-Compartments of *Drosophila* Secondary Cells Contain Intraluminal Vesicles with Specific Cargos. (A) Schematics illustrate the fly accessory glands and associated secondary cells, with their exosome forming compartments. First panel shows male fruit fly and its accessory glands. Boxed region is enlarged in second panel, revealing secondary cells (SCs; black dots), with red line indicating the plane of section through the AG lumen, which is used to generate the transverse view through an SC and surrounding main cells within the epithelial layer in the third panel. The unusually large, acidic late endosomes and lysosomes (Rab7-positive; magenta) and non-acidic compartments, characteristic of SCs, which we demonstrate here contain intraluminal vesicles (ILVs; some of which are Rab11-positive; yellow), are labelled. The red line in the third panel shows the basal SC plane of section shown in the fourth panel and images in this figure. Panels B-E show basal views through living SCs, with dashed white circles approximating the outline of a single SC, and acidic compartments marked by the vital dye LysoTracker^®^ Red (magenta). In merge images, a single non-acidic (B, C, E) and acidic (D) compartment containing intraluminal vesicles (ILVs) is boxed and magnified in the right panel (Zoom). ILVs appear as membrane-delineated vesicles, using super-resolution 3D-structured illumination (3D-SIM) microscopy for the brighter overexpressed GFP-tagged constructs (yellow; B and E). However, ILVs appear only as puncta, using lower resolution wide-field microscopy for the fainter endogenously expressed YFP-tagged Rab GTPases (yellow; C and D). (B) 3D-SIM image of SC expressing a GFP-tagged version of human CD63 (CD63-GFP). CD63-GFP expression is apparent on the limiting membranes of non-acidic compartments and their ILVs, and also the limiting membranes of the enlarged acidic compartments (also seen in Figure S2A). Arrow highlights CD63-GFP-marked ILVs (Zoom). Many more non-acidic compartment ILVs are apparent in a complete Z-stack of a non-acidic compartment (see Supplementary Movie S1). (C) Wide-field fluorescence image of an SC expressing a *YFP-Rab11* gene trap. YFP-Rab11 marks the limiting membranes of most non-acidic compartments and internal puncta (arrow in Zoom), but not the surface of acidic compartments (Figure S2B). (D) Wide-field fluorescence image of SC expressing a *YFP-Rab7* gene trap. YFP-Rab7 marks the limiting membranes of acidic compartments and internal puncta (arrow in Zoom). Enlarged acidic compartments are also present in adjacent main cells. (E) 3D-SIM image of SC expressing a GFP-tagged version of Breathless (Btl-GFP), the fly FGFR homologue. Btl-GFP marks the limiting membranes of non-acidic compartments and their ILVs (arrow in Zoom), but not the surface of acidic compartments (also seen in Figure S2C). Images from six-day-old male flies shifted to 29°C at eclosion. This induces GAL4/UAS-dependent SC transgene expression in (B) and (E). The genotypes of flies carrying multiple transgenes are: *w; P[w^+^, UAS-CD63-GFP] P[w^+^, tub-GAL80^ts^]/+*; *dsx-GAL4/+* (B); *w; P[w^+^, tub-GAL80^ts^]/+*; *dsx-GAL4/P[w^+^, UAS-btl-GFP]* (E). Scale bar in B-E (5 µm) and in B-E, Zoom (2 µm). See also Supplementary Figures S1, S2 and Movies S1, S2.

Here we present evidence that a specific exosome subtype is made in the Rab11-positive compartments of SCs and in human tumour cell recycling endosomal MVBs labelled by Rab11 family members. Furthermore, preferential release of Rab11a-marked exosomes from these compartments is triggered by depleting cancer cells of exogenous glutamine. This Rab11a-secretory switch is reproduced by reducing growth factor-regulated Akt and amino acid-sensitive mechanistic (formerly mammalian) Target of Rapamycin Complex 1 (mTORC1; Dibble and Cantley, 2015) signalling. We show that these exosomes have distinct cargos and unique *in vitro* activities. Since they also alter tumour cell growth and vessel formation in xenografts *in vivo*, we propose that they contribute to adaptive responses to metabolic stress.

## RESULTS

### Rab11-Labelled Multivesicular Bodies Make Exosomes via an ESCRT-Dependent Mechanism in *Drosophila* Secondary Cells

To study exosome biogenesis in *Drosophila* SCs, we overexpressed the human exosome marker CD63-GFP. It labels the limiting membranes of large Rab7-positive acidic late endosomes and lysosomes (LELs) in SCs and of approximately ten non-acidic compartments, containing a central, protein-rich, dense-core granule (DCG; Figures 1A, 1B, S1A), which are Rab11-positive (Redhai et al., 2016; Prince et al., 2019). Using super-resolution 3D-SIM, a few fluorescent ILVs were observed inside LELs, although most GFP fluorescence is quenched by the acidic microenvironment (Redhai et al., 2016). Clusters of CD63-GFP-positive ILVs were also observed within the Rab11-positive compartments (Figure 1B; Movie S1), the majority of which were of exosome size (Figure S1F). ILVs were also seen in EM micrographs of the non-acidic and acidic compartments in non-transgenic flies (Figure S1G).

To analyse ILV formation in these Rab11 compartments further, we studied SCs from flies expressing YFP-Rab11 from the endogenous *Rab11* locus, a so-called ‘gene trap’ (Dunst et al., 2015). We imaged this marker using wide-field deconvolution fluorescence microscopy, which resolves membrane-bound ILVs as puncta, since YFP-Rab11’s low fluorescence intensity relative to CD63-GFP was not amenable to super-resolution imaging. The compartmental organisation of SCs expressing *YFP-Rab11* does not appear significantly perturbed, in contrast to SCs expressing CD63-GFP (Figure S1H-J; Redhai et al., 2016). For example, numbers of large non-acidic compartments are unaffected compared to wild-type flies, whereas CD63-GFP expression roughly doubles their number (Figure S1H; Redhai et al., 2016). Fluorescent puncta were seen inside YFP-Rab11-positive SC compartments (Figure 1C), but fewer than with CD63-GFP (Figure 1B). Moreover, unlike CD63-GFP (Figure S1B), the Rab11 fusion protein did not traffic to the plasma membrane (Figure S1C, Movie S2). Sporadic YFP-Rab11-positive puncta were observed in the AG lumen, both in the gene trap line (Figure S1C) and when YFP-Rab11 was specifically overexpressed in SCs (Figure S1D). In contrast, a YFP-Rab7 gene trap fusion protein (Dunst et al., 2015) primarily trafficked to acidic LELs (Figure 1D) and marked very few puncta in the AG lumen. We conclude that Rab11-labelled exosomes are formed at low levels in Rab11 SC compartments and secreted.

In searching for other markers of these alternative exosomes, we found that an overexpressed GFP-tagged form of Breathless (Btl; a fly homologue of the human transmembrane FGF receptor), which is normally expressed in SCs (Figure S1E), trafficked on to ILVs in Rab11-compartments (Figure 1E). Its expression did not affect large non-acidic compartments in SCs and had only a minor effect on acidic compartment number (Figure S1H-J). In some SCs, Btl-GFP, like YFP-Rab11, was found in the lumen of an LEL, but, unlike CD63-GFP, these markers were rarely, if ever, observed on the LEL limiting membrane (Figure S2), presumably because they reach LELs by sporadic fusion to Rab11-compartments (Corrigan et al., 2014) and not by endocytic trafficking. As with YFP-Rab11, transmembrane Btl-GFP protein was secreted in puncta into the AG lumen (Figure 2A). We, therefore, conclude that Rab11 and Btl are selective membrane-associated markers for exosomes generated in Rab11-compartments of SCs, which we describe as ‘Rab11-exosomes’.

**Figure 2.**
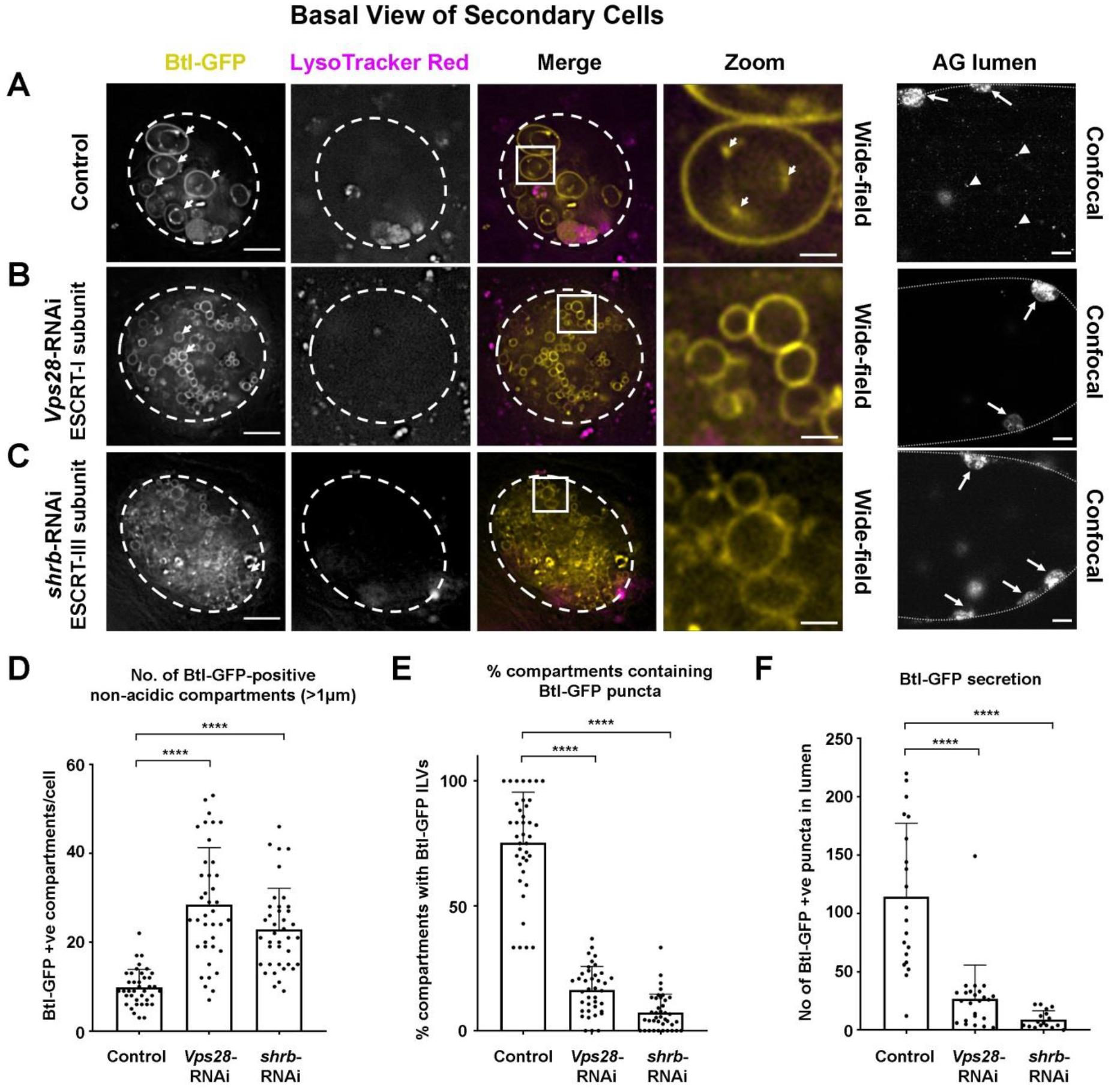
Exosome Biogenesis in Rab11-Compartments of *Drosophila* Secondary Cells is ESCRT-Dependent. Panels A-C show wide-field fluorescence images of basal views through living SCs expressing a GFP-tagged form of the fly FGFR homologue, Breathless (Btl-GFP; yellow) in non-acidic compartments, with SC outline approximated by dashed white circles. Acidic compartments are marked by LysoTracker Red^®^ (magenta). Boxed non-acidic compartments in merge images are magnified in A-C, Zoom. Lower magnification confocal transverse images of fixed accessory gland (AG) lumens are on the right-hand side with dotted lines indicating the AG epithelial layer, which contains at least one fluorescent SC (highlighted by arrows). (A) SC with no RNAi construct expressed (control) and AG lumen from same genotype. Btl-GFP-positive ILV membranes are apparent inside compartments (arrowheads in Zoom) and as secreted puncta (AG lumen; arrowheads). (B) SC also expressing RNAi construct targeting ESCRT-I subunit, *Vps28*, and AG lumen from same genotype. Btl-GFP-positive ILVs (Zoom) and secreted puncta (AG lumen) are strongly reduced. (C) SC also expressing RNAi construct targeting ESCRT-III subunit, *shrb*, and AG lumen from same genotype. Btl-GFP-positive ILVs (Zoom) and secreted puncta are strongly reduced. (D) Bar chart showing the number of large (diameter greater than one micrometre) non-acidic Btl-GFP-positive compartments per SC. Data from 39 SCs (three per gland) are shown. (E) Bar chart showing percentage of Btl-GFP compartments containing Btl-GFP-positive ILVs in control and *ESCRT* knockdown SCs. Data from 39 SCs (three per gland) are shown. (F) Bar chart showing the total number of Btl-GFP fluorescent puncta in ten Z-planes from AG lumen following *ESCRT* knockdown in SCs, compared to controls without knockdown. Data from at least 17 AG lumens per condition are shown. All data are from six-day-old male flies shifted to 29°C at eclosion to induce expression of transgenes. Genotypes are: *w; P[w^+^, tub-GAL80^ts^]/+*; *dsx-GAL4/P[w^+^, UAS-btl-GFP]* with no knockdown construct (A), UAS-*Vps28*-RNAi (v31894; B) or UAS-*shrb*-RNAi (v106823; C). Scale bars in A-C (5 µm); in A-C Zoom (1 µm); in A-C, AG lumen (20 µm). Data were analysed by one-way ANOVA. **P < 0.01 relative to control. See also Figures S2, S3 and Supplementary Movie S3.

To test whether biogenesis of these exosomes is ESCRT-dependent, the temperature-inducible GAL4/GAL80^ts^/UAS system was used to knock down two *ESCRT*s implicated in mammalian and *Drosophila* exosome secretion (Matusek et al., 2014), namely *Vps28* (an ESCRT-I subunit) and *shrub* (*shrb*; the fly orthologue of mammalian *Chmp4a-*c, encoding an ESCRT-III subunit) in adult SCs. Both treatments affected the number and size of large non-acidic compartments in SCs expressing Btl-GFP (Figure 2), YFP-Rab11 and CD63-GFP (Figure S3). In Btl-GFP-expressing SCs, the number of non-acidic Rab11-compartments with diameter greater than one micrometre increased in both knockdowns (Figure 2A-D), but analysis of *Rab11* gene trap males indicated that although drastically reduced in size, many of these compartments retained Rab11 identity (Figure S3A-C). In both *ESCRT* knockdowns, very few compartments contained Btl-GFP puncta and the number of secreted Btl-GFP exosomes in the AG lumen was greatly reduced (Figure 2A-C, E, F). A similar inhibition of ILV formation in non-acidic compartments and exosome secretion was produced by *ESCRT* knockdown in CD63-GFP-expressing SCs (Figure S3D-F). To test whether ESCRT proteins associate with Rab11-compartments, a Shrb-GFP fusion protein (Sweeney et al., 2006) was transiently expressed in SCs. It accumulated in sub-domains and puncta at the surface of large non-acidic and acidic compartments (Figure S3G; Movie S3), consistent with our finding that *shrb* plays a role in generating ILVs in Rab11-compartments, in addition to the well-established LEL compartments. We conclude that the ESCRTs, Vps28 and Shrb, are required for ILV formation in Rab11-compartments *in vivo*, and that these ILVs are normally secreted as exosomes loaded with specific cargos.

### Rab11a-Labelled Multivesicular Bodies Also Generate Intraluminal Vesicles in HCT116 Colorectal Cancer Cells

In human cells, Rab11a, one of the two Rab11 isoforms, which primarily associates with recycling endosomes, has been reported to be associated with EVs (Keerthikumar et al., 2016). It also regulates exosome secretion (Savina et al., 2002). In HCT116 colorectal cancer cell (CRC) cells, which have clustered endosomes, Rab11a compartments are distinct from those marked by the LEL protein LAMP2 (Figure 3A) and CD63 (Figure 3B), which co-localises with the LEL marker, LAMP1 (Figure 3C). This contrasts with CD63-GFP in SCs, where some overexpressed protein enters Rab11 compartments (Figure 1B). By overexpressing GFP-Rab11a in these cells, which induces clustering of Rab11a-positive compartments, we were able to use super-resolution 3D-SIM microscopy to detect internalised GFP in these compartments (Figure 3D), suggesting that Rab11a is incorporated into ILVs. To enlarge endosomal compartments, a GFP-tagged, constitutively active form of early endosomal Rab5 was expressed, which inhibits maturation of recycling and late endosomes, leading to the formation of enlarged immature endosomal compartments (Figure 3E). The endosomal Rab GTPases, Rab11a and Rab7, were both observed in sub-domains at the surface of these Rab5-positive compartments and also on spatially distinct internal puncta (Figure 3E), demonstrating that the endosomal system can generate ILVs carrying either of these Rab signatures. We conclude that HCT116 cells contain both late and recycling endosomal MVBs and, as apparent in *Drosophila* SCs, the ILVs produced within them carry different cargos, including specific Rab GTPases.

**Figure 3.**
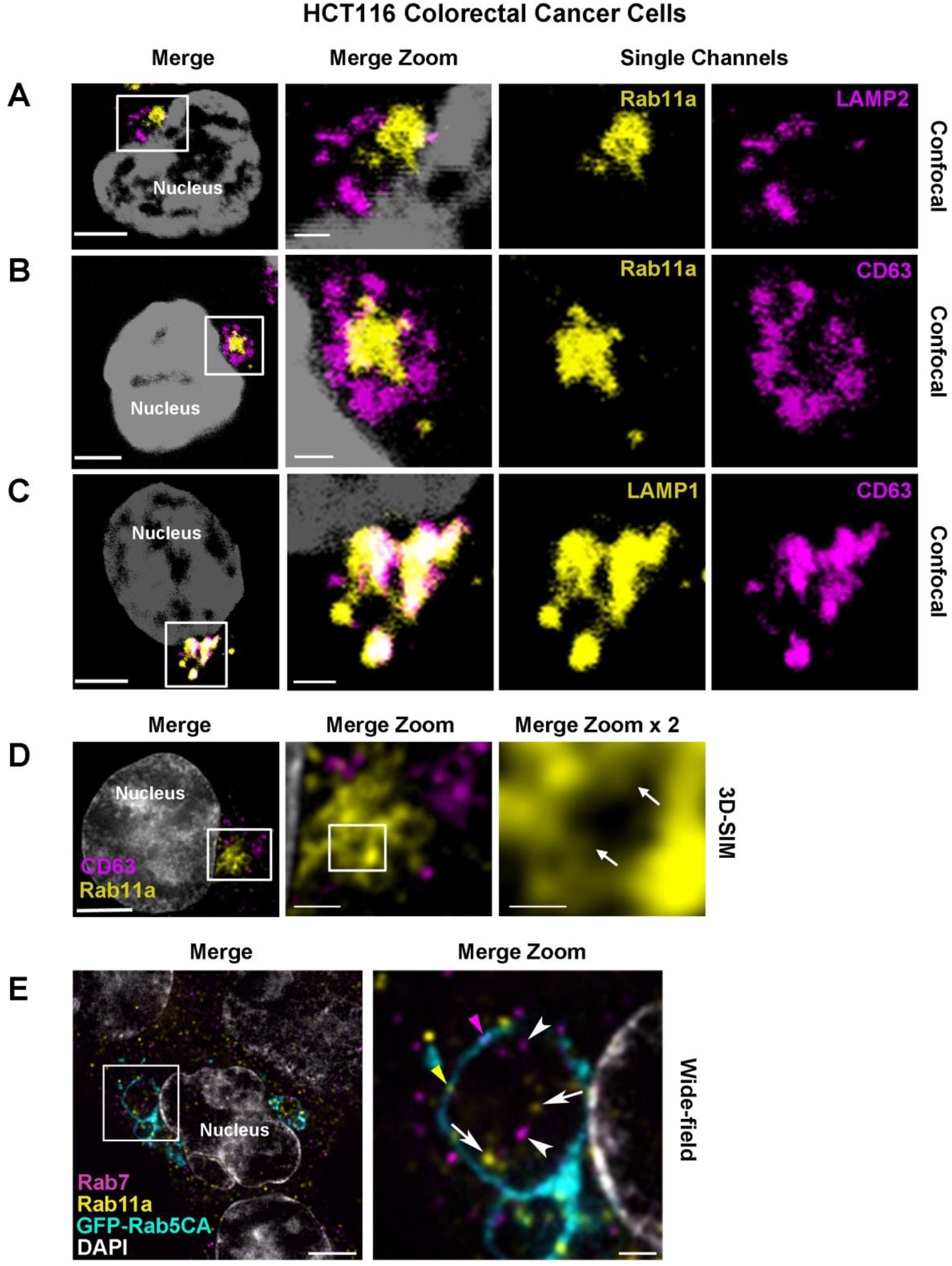
Rab11a Labels a Distinct Subset of Multivesicular Bodies and Their Intraluminal Vesicles in HCT116 Colorectal Cancer Cells. Panels A-C show confocal images of fixed HCT116 cells, with boxed regions enlarged to the right. DAPI (grey) marks nucleus. (A) Rab11a (yellow) is located in compartments distinct from the late endosomal and lysosomal marker, LAMP2 (magenta). (B) Rab11a (yellow) is located in compartments distinct from CD63 (magenta). (C) CD63 (magenta) predominantly co-localises with the late endosomal and lysosomal marker, LAMP1 (yellow). (D) Super-resolution 3D-SIM image of fixed HCT116 cell expressing GFP-Rab11a (yellow), and stained with CD63 (magenta). DAPI (grey) marks nucleus. Boxed Rab11a-positive compartments, which frequently cluster, are magnified in Merge Zoom. This panel is further magnified in Merge Zoom x 2, revealing GFP-Rab11a (arrows in right panel) inside compartments. (E) Wide-field fluorescence image of fixed HCT116 cells, stained with Rab11a (yellow) and Rab7 (magenta), expressing constitutively active GFP-tagged Rab5 (GFP-Rab5CA; cyan), which stalls endosomal maturation and produces enlarged Rab5-positive endosomes. One of these is boxed in the Merge and magnified in Merge Zoom, revealing internal puncta marked by Rab11a (arrows) and Rab7 (arrowheads) and limiting membrane subdomains of Rab11a (yellow arrowhead) and Rab7 (magenta arrowhead). DAPI (grey) marks nuclei. Scale bars in A-E (5 µm), in Merge Zoom (1 µm), and in Merge Zoom x 2 (0.5 µm).

### Glutamine Depletion of HCT116 Cells Induces a Switch to Secretion of Rab11a-Exosomes with Distinct Cargos

To explore whether HCT116 CRC cells secrete Rab11a-positive exosomes, we collected small EVs for 24 h from cells cultured in serum-free conditions, but supplemented with insulin. This maintained growth factor signalling, as assessed by phosphorylation of mTORC1 downstream read-outs, 4E-Binding Protein 1 (4E-BP1) and S6 (2.00 mM glutamine in Figure 4A). EVs isolated by ultracentrifugation were enriched for proteins that preferentially associate with exosomes (Figure 4B, S4D), namely, the cytosolic adaptor protein Syntenin-1 (Syn-1), the ESCRT-I component Tsg101 and the tetraspanins CD81 and CD63 (Kowal et al., 2016). Low levels of Rab11a were also present, but not ER, Golgi or early endosome markers.

**Figure 4.**
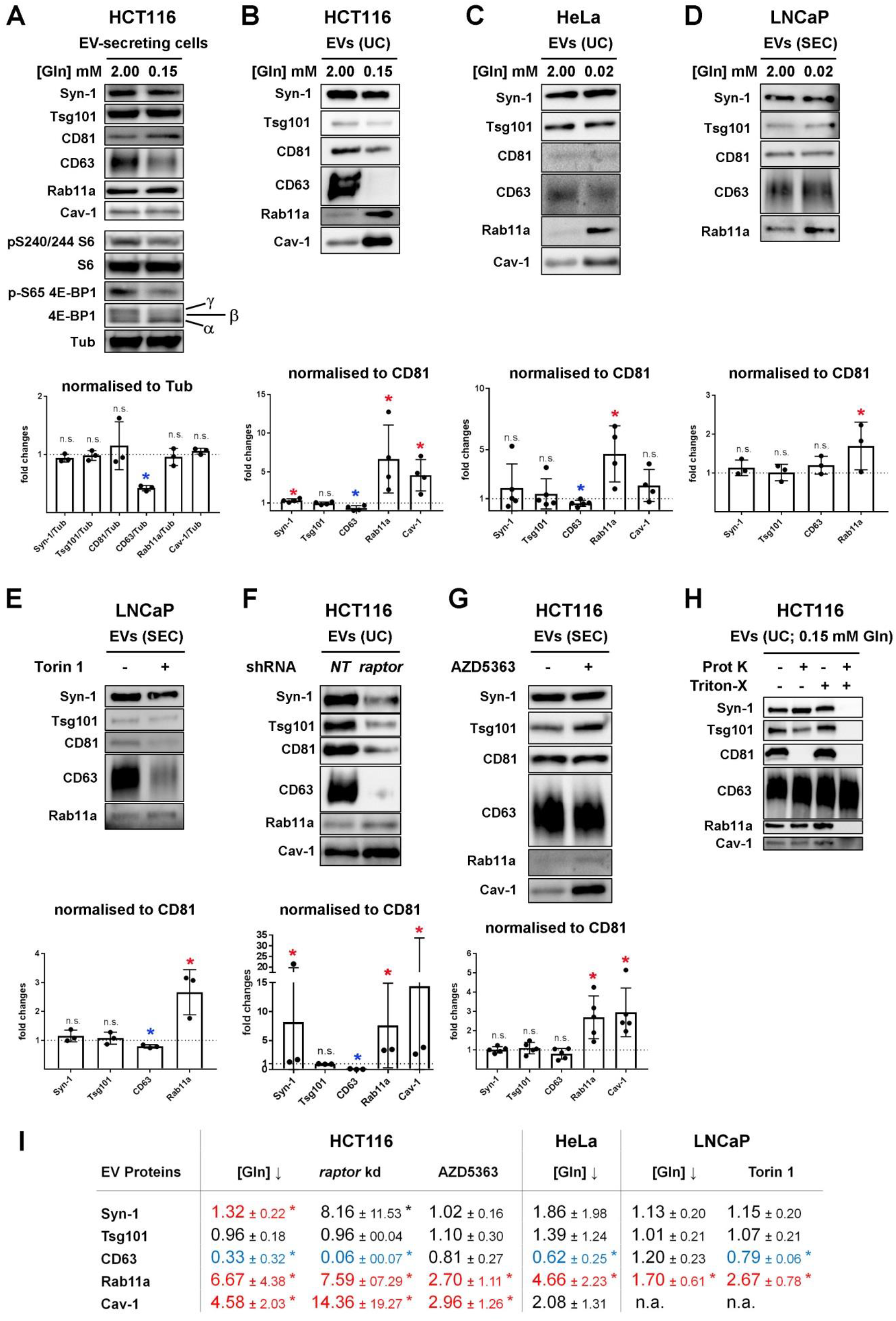
Reduction in Extracellular Glutamine or Akt/mTORC1 Signalling Induces a Switch to Rab11a-Positive Exosome Secretion in Three Human Cancer Cell Lines. (A) Western blot analysis of putative exosome markers in lysates from HCT116 cells cultured in glutamine-replete (2.00 mM) and glutamine-depleted (0.15 mM) medium for 24 h. Gel was loaded with equal amounts of protein (total cell lysate protein was reduced by 19 ± 4 % after glutamine depletion). The activity of mTORC1 was assessed via phosphorylation of S6 and 4E-BP1, using phospho-specific antibodies and a pan-4E-BP1 antibody, where the hyper-phosphorylated form of 4E-BP1 produced by mTORC1 gives the slowest migrating γ band. Bar chart shows the abundance of putative exosome proteins relative to tubulin in these lysates Panels B-H show western blot analyses of EV preparations with loading normalised to cell lysate protein levels. In the bar charts, the levels of putative exosome proteins are normalised to CD81. (B) EVs isolated by ultracentrifugation (UC) of medium from HCT116 cells cultured in glutamine-replete and -depleted conditions for 24 h. (C) EVs isolated by UC of medium from HeLa cells cultured in glutamine-replete (2.00 mM) and -depleted (0.02 mM) conditions for 24 h. (D) EVs isolated by size-exclusion chromatography (SEC; fractions two to seven) from LNCaP cells cultured in glutamine-replete (2.00 mM) and -depleted (0.02 mM) conditions for 24 h. (E) EVs isolated by SEC (fractions two to seven) from LNCaP cells cultured in the presence or absence of 120 nM Torin 1 for 24 h. (F) EVs isolated by UC from HCT116 cells cultured for 24 h following three days of *raptor* or non-targeting (*NT*) shRNA knockdown. (G) EVs isolated by SEC from HCT116 cells cultured in the presence or absence of 3 µM Akt inhibitor AZD5363 for 24 h. (H) EVs isolated by UC under glutamine-depleted conditions as in (B) and then subjected to Proteinase K (Prot K) digestion in the presence or absence of Triton^®^ X-100 (Triton-X). Only membrane-associated CD81 is digested in the absence of Triton^®^ X-100, while CD63 is resistant to digestion, even in the presence of Triton^®^ X-100. (I) Table summarising the EV protein analyses in panels B to G (normalised to CD81 levels). Significantly decreased levels are in blue and increased levels are in red. Bar charts derived from at least three independent experiments and analysed by the Kruskal-Wallis test: *P < 0.05; n.s. = not significant. See also Figures S4, S5.

Glutamine is a major metabolic substrate in HCT116 cells (Jiang et al., 2013). We have previously shown that nutrient stress induced by glutamine depletion inhibits a rapamycin-resistant form of nutrient-sensitive mTORC1 (Fan et al., 2016). We investigated whether cells respond to such treatment by altering their EV secretion, as they do in hypoxic stress (Kucharzewska et al., 2013). As expected, glutamine depletion reduced cellular levels of hyper-phosphorylated 4E-Binding Protein 1 (4E-BP1) and phosphorylated S6 (Figure 4A, S4A) over the 24 h EV collection period. Cellular expression of all exosome proteins was unaffected by glutamine depletion, except for CD63, which was reduced (Figure 4A). There were comparably low levels of cell death in secreting cells under both glutamine-depleted (10.2 ± 1.5%) and -replete (11.5 ± 2.5%; n=3) conditions, as measured by Trypan Blue staining.

Nanoparticle Tracking Analysis (NTA) of EV preparations (Dragovic et al., 2011) revealed that EV numbers (when normalised to cell lysate protein levels) were not significantly changed compared to control values after glutamine depletion, and EV size distribution was unaltered (Figure S4B, S4E), a finding supported by transmission electron microscopic analysis (Figure S4C).

However, we observed that glutamine depletion of HCT116 cells altered the secretion of specific exosome markers (Figure 4B, I). While exosome-associated Syn-1, Tsg101 and CD81 were only slightly decreased after glutamine depletion, when normalised to cell lysate protein levels (a proxy for cell number), late endosomal CD63, which is reported to mark a subset of exosomes in small EV preparations (Kowal et al., 2016), was much more strongly reduced (Figure S4E). By contrast, Rab11a was increased by several-fold (Figure 4B, S4E). Hypoxic stress increases levels of the cytosolic lipid raft-associated Caveolin-1 (Cav-1; Kucharzewska et al., 2013) in EVs. This protein was detected in EV preparations from HCT116 cells and its secretion was also enhanced by glutamine depletion (Figure S4E). While no marker is known to distribute evenly across all exosome subtypes, when CD81, Tsg101 or Syn-1 were employed to normalise western blots of EVs (Figure 4B, S4E), levels of CD63 always fell significantly following glutamine depletion, which may be partly explained by the reduction in cellular CD63 (Figure 4A), but Rab11a and Cav-1 levels were strongly increased. This mirrored our findings when protein levels in EV-secreting cells were used for normalisation. To standardise our analysis, levels of endosomal CD81 from glutamine-replete and-depleted conditions were used for normalisation of EV preparations in subsequent experiments (Figure 4B, I). The equivalent Rab11a-switch was also observed in EVs isolated by size-exclusion chromatography (SEC; Figure S4F, S4G), an alternative method for EV isolation. We conclude that following glutamine depletion, the endosomal origin of exosomes secreted from CRC cells is altered, so that more exosomes are released from Rab11a compartments.

To confirm that endosomal Rab11a is a *bona fide* internal exosome marker and not a contaminant in the ultracentrifugation procedure, EVs induced by glutamine depletion were subjected to Proteinase K digestion in the absence and presence of the detergent Triton X-100. While proteins partially exposed on the exosome surface like CD81 were affected by both treatments, Rab11a behaved like other internal markers, such as Syn-1, and was only digested in the presence of detergent (Figure 4H).

### Glutamine Depletion and Inhibition of Akt/mTORC1 Signalling Induces an ‘Exosome Switch’ in Cancer Cell Lines of Different Origins

We tested the effect of glutamine depletion on EV secretion by other cancer cell lines. Culturing HeLa cervical cancer cells in glutamine-depleted medium, which reduced phosphorylation of S6 and 4E-BP1 (Figure S5A, S5H), slightly increased EV number (Figure S5A’, S5H), but greatly increased Rab11a secretion relative to all other markers (Figure 4C, I; an apparent increase in EV levels of Cav-1 did not reach significance), while cellular levels of Rab11a were unaffected (Figure S5A). In the prostate cancer cell line, LNCaP, depletion of extracellular glutamine, which also reduced S6 and 4E-BP1 phosphorylation (Figure S5B, S5H), led to increased levels of Rab11a in EVs relative to CD81, while CD63 and EV number were not significantly changed (Figure 4D, I, S5B’, S5H). Cav-1 was not detectable in this cell line. We conclude that HeLa and LNCaP cells, like HCT116 cells, respond to glutamine depletion by increasing secretion of exosomes containing Rab11a.

Since glutamine depletion reduces mTORC1 activity, we tested whether blocking mTORC1 with the mTOR-specific inhibitor Torin 1 (Thoreen et al., 2009) could also induce an exosome switch. In LNCaP cells, Torin 1 treatment slightly elevated the number of EVs produced, but reduced CD63 and strongly increased Rab11a in EVs relative to CD81 (Figure 4E, I), without affecting cellular levels of these proteins (Figure S5C, S5C’, S5H). However, in other cell lines like HCT116, the mTORC1 inhibitors, Torin1 and rapamycin, had a much stronger inhibitory effect on mTORC1 activity and particularly S6 phosphorylation when compared to glutamine depletion (Figure S5D, S5E, S5I), and this was associated with a large reduction in EV numbers (Figure S5D”, S5E”, S5H). Secretion of all exosome markers was significantly reduced (Figure S5D’, S5E’), suggesting a general shut-down in exosome release.

A more modest mTORC1 inhibition in HCT116 cells was induced by knocking down *raptor*, which encodes a core mTORC1 component. EVs were collected for 24 h after three days of knockdown. Under these conditions, cellular levels of Syn-1, Tsg101 and CD63 were all decreased (Figure S5F) and EV number was also reduced (Figure S5F’, S5H), but residual phosphorylation of S6 and 4E-BP1 was maintained (Figure S5F, S5H). This treatment induced a reduction in several secreted exosome proteins, with a stronger reduction in CD63. However, there was an increase in Rab11a and Cav-1 relative to CD81 (Figure 4F, I), consistent with induction of an mTORC1-regulated switch in the balance of exosome secretion from late endosomal to Rab11a-compartments.

Since mTORC1 activity is controlled by growth factor signalling, we also tested the effect of the Akt inhibitor AZD5363 on exosome secretion by HCT116 cells. A switch in exosome secretion was again observed (Figure 4G, I), most notably involving an increase in Rab11a and Cav-1 in EVs, in the absence of similar changes in cellular levels of these proteins or EV number (Figures S5G, S5G’, S5H). In addition to blocking phosphorylation of the Akt target PRAS40, this treatment partially inhibited phosphorylation of mTORC1 targets, S6 and 4E-BP1 (Figure S5G, S5H), which might account for the switch. Overall, our data from CRC, cervical and prostate cancer cell lines lead us to conclude that glutamine depletion or inhibition of Akt/mTORC1 signalling induces a switch towards secretion of vesicles from recycling endosomal compartments, increasing the levels of Rab11a-exosomes by several fold. However, strong mTORC1 blockade can suppress all exosome secretion, so this switch is no longer apparent.

### Glutamine-Depletion-Induced Extracellular Vesicles Have Altered Activities That Are Dependent on Rab11a

Since EVs secreted by glutamine-depleted HCT116 cells have altered cargos, we tested whether this affected their bioactivity. When EVs from glutamine-replete and -depleted cells were prepared by ultracentrifugation, then mixed with naïve recipient HCT116 cells, 30 min prior to plating, the stress-induced EVs selectively stimulated cell growth under low serum conditions (Figure 5A). Cell proliferation was enhanced by this treatment (Figure S4H), but rates of apoptosis were unaffected (Figure S4I). The same activity was reproduced with EVs isolated by SEC, and shown to be dose-dependent (Figures 5A, S4J), so both of these methods were employed in subsequent functional assays.

**Figure 5.**
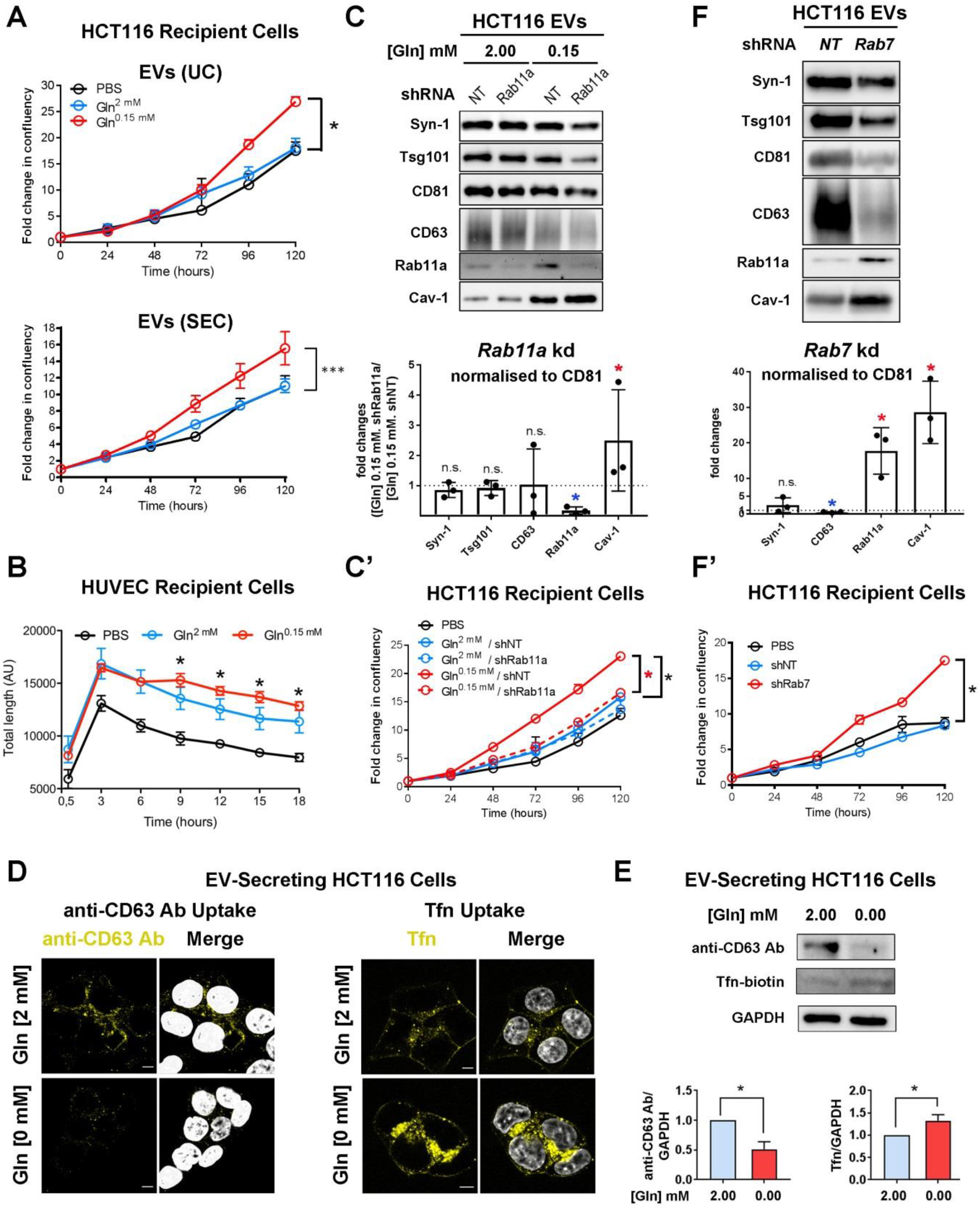
Rab11a-Dependent Exosome Secretion is Induced by Altered Endosomal Trafficking and Promotes Recipient Tumour Cell Growth. (A) Growth curves for HCT116 recipient cells in reduced (1%) serum conditions following 30 min pre-incubation with EVs isolated by UC (10^4^ per cell; top) or by SEC (4 x 10^3^ per cell; bottom) from glutamine-replete and -depleted HCT116 cells or with vehicle (PBS). (B) Cumulative tube length for HUVEC recipient cells following treatment with 10^4^ EVs isolated by UC from glutamine-replete and -depleted HCT116 cells or with vehicle (PBS). Both EV preparations promote tubulation, but the network is more stable with EVs from glutamine-depleted HCT116 cells. (C) Western blot analysis of EVs isolated by UC from HCT116 cells cultured in glutamine-replete and -depleted medium for 24 h, following transduction with a *Rab11a* or control non-targeting (NT) shRNA knockdown construct over previous two days. Bar chart shows change in putative exosome proteins in EVs secreted from *Rab11a* knockdown cells relative to NT-treated cells in glutamine-depleted conditions, following initial normalisation to CD81. (C’) Growth curves are for HCT116 recipient cells pre-treated with EVs isolated as in (C) or with vehicle (PBS). (D) Left hand group of four images show cells grown in glutamine-replete (2.00 mM) and glutamine-starved (0.00 mM) conditions for 24 h, then incubated with an anti-CD63 antibody (yellow) for 30 min at 4°C, washed with PBS, chased at 37°C for 30 min, then fixed, immunostained and imaged. Right hand group of four images show cells grown in glutamine-replete and glutamine-starved conditions for 24 h, incubated with Tfn-Alexa Fluor^®^ 488 (yellow) for 30 min at 4°C, shifted to 37°C for 30 min, then fixed and imaged. (E) Western blot showing levels of anti-CD63 heavy chain and biotin-conjugated Tfn in HCT116 cells cultured for 24 h in glutamine-replete or -starved conditions, incubated for 30 min in medium containing these molecules at 4°C, then chased at 37°C for 30 min (the wash step was not performed for Tfn in D, to reduce loss due to rapid recycling of Tfn). (F) Western blot analysis of EVs isolated by UC from HCT116 cells in glutamine-replete medium, transduced with a *Rab7* or non-targeting (NT) control shRNA knockdown construct. Bar charts show changes in putative exosome proteins in EVs isolated by ultracentrifugation, following normalisation to CD81. (F’) Growth curves are for HCT116 recipient cells pre-treated with EVs isolated as in (F) or with vehicle (PBS). Growth curves were reproduced in three independent experiments. Scale bars in D (5 µm). Bar charts derived from three independent experiments: *P < 0.05. See also Figures S4 and S6.

To test for effects of glutamine-depletion-induced EVs on stromal cells, EV preparations were also added to human umbilical vein endothelial cells (HUVECs), which were then plated in Matrigel^®^ to assess their ability to form and maintain a tubular network. EVs isolated under glutamine-replete and -depleted conditions increased network formation compared to PBS-treated controls (Figure 5B). Network maintenance was, however, enhanced by EVs from glutamine-depleted cells, suggesting that these EVs are better at supporting a stable capillary network. Therefore, following glutamine depletion, HCT116 cells secrete EVs enriched in exosomes from Rab11a-positive endosomes that display novel or enhanced activities.

To test whether trafficking through the Rab11a-dependent recycling endosomal pathway was essential for the increased growth induced by EVs from glutamine-depleted cells (Figure 5A), *Rab11a* was knocked down in secreting cells. This reduced the number of EVs secreted by approximately 40% (Figure S6A’), but had no effect on mTORC1 activity or cellular levels of exosome proteins (Figure S6A). Exosome markers Syn-1, Tsg101 and CD63 were unaltered in EVs compared to controls and Cav-1 secretion was increased (Figure 5C), suggesting that Rab11a-compartments are not the only route for secretion of these molecules. Knockdown did, however, strongly suppress the enhanced growth-promoting activity of EVs produced by glutamine-depleted cells (Figure 5C’). We conclude that this activity is likely to involve Rab11a-dependent exosome secretion.

### Altered Endosomal Pathway Trafficking Induces an Exosome Switch

We reasoned that changes in endosomal trafficking might be involved in switching the balance of exosome secretion to favour Rab11a-exosomes following glutamine depletion. To test this, the internalisation of molecules that specifically traffic from the plasma membrane either to LELs or to Rab11a-compartments was assessed under both glutamine-replete and - depleted conditions. While CD63 accumulates in LEL compartments (Figure 3C), Transferrin (Tfn) is returned to the plasma membrane after endocytosis, via the rapid recycling pathway and Rab11a-positive recycling endosomes (Mayle et al., 2012). Consistent with this, internalised Alexa-488-conjugated Tfn partly co-localised with Rab11a in HCT116 cells (Figure S6B), but not with the LEL marker LAMP2 (Figure S6C). When HCT116 cells were depleted of glutamine, uptake of an anti-CD63 antibody, assessed by confocal imaging and western analysis, was significantly reduced (Figure 5D, E), consistent with the reduced levels of CD63 in these cells. By contrast, uptake of fluorescent (Figure 5D) and biotin-conjugated (Figure 5E) Tfn was elevated, demonstrating that flux through the recycling endosomal pathway is increased in response to glutamine depletion.

Since altered endosomal trafficking is associated with a glutamine-regulated switch in exosome secretion, we hypothesised that a similar switch might be induced in HCT116 cells by inhibiting traffic through the late endosomal pathway. Knockdown of *Rab7* substantially reduced EV secretion, decreased CD63 and several other exosome markers in EV preparations under glutamine-replete conditions, but increased Rab11a levels in EVs, without having a major effect on marker expression levels in cells (Figure 5F, S6D and D’; cellular levels of Rab11a were in fact slightly reduced). The resulting EVs stimulated growth of serum-depleted HCT116 cells (Figure 5F’), demonstrating that altering the balance of late and recycling endosomal trafficking can induce changes analogous to the switch in exosome secretion caused by reducing glutamine.

### EVs Secreted from Glutamine-Depleted Cells Induce ERK-Dependent Growth in HCT116 Recipient Cells and Are Blocked by an Anti-Amphiregulin Antibody

Since EVs from glutamine-depleted HCT116 cells induce growth in recipient cells, we investigated whether growth factor signalling might be involved. Phosphorylation of the MAPK ERK was enhanced when serum-starved HCT116 cells were incubated for 30 min with EV preparations from glutamine-depleted cells (Figure 6A). Indeed, treating recipient HCT116 cells with the ERK inhibitor SCH772984 for the first 24 h of culture after EV addition completely blocked the extra growth induced by these vesicles (Figure 6B), demonstrating that elevated ERK signalling is critically involved in growth stimulation.

**Figure 6.**
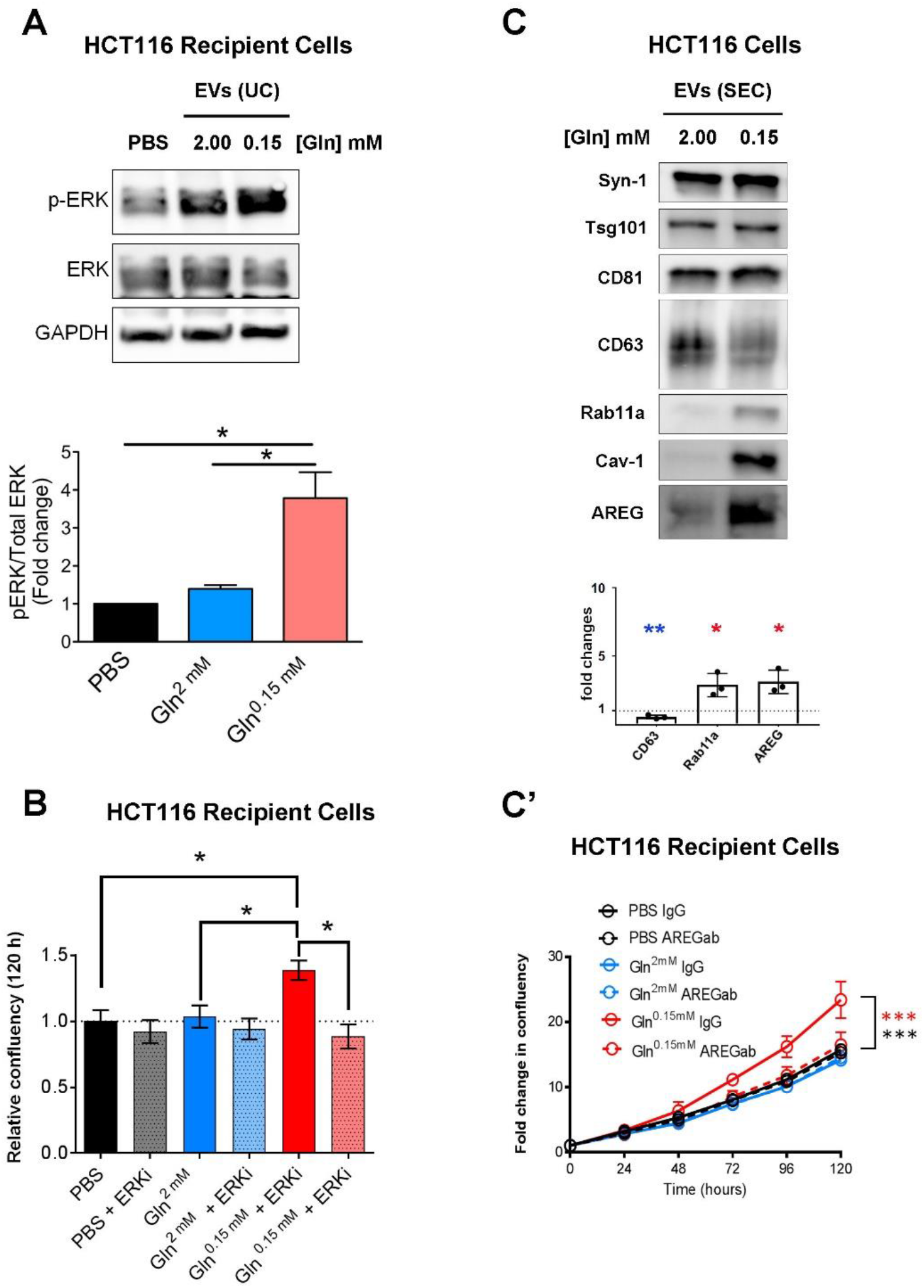
EVs Induced by Glutamine Depletion Promote ERK-and Amphiregulin-Dependent Growth in HCT116 Recipient Cells. (A) Western blot analysis of ERK phosphorylation (p-ERK) in recipient serum-deprived HCT116 cells pre-treated with EVs isolated by UC from glutamine-replete or -depleted HCT116 cells or vehicle (PBS). Bar chart shows ratio of p-ERK to ERK levels from triplicate independent experiments. (B) Bar chart shows HCT116 recipient cell growth over 120 h, after pre-treatment with EV preparations isolated as in (A) or with PBS, and then incubation for the first 24 h of culture in the presence or absence of the ERK inhibitor SCH772984 (1.00 µM). (C) Western blot analysis showing levels of the EGFR ligand, Amphiregulin (AREG) in EVs isolated by SEC of medium from HCT116 cells cultured in glutamine-replete (2.00 mM) or - depleted (0.15 mM) conditions for 24 h. Loading was normalised to cell lysate protein levels. AREG’s molecular weight (approximately 26-28 kDa) suggests it is in its membrane-associated form. In the bar chart below, the levels of putative exosome proteins were normalised to CD81. (C’) Growth curves in low (1%) serum are for HCT116 recipient cells pre-treated with the EV preparations [isolated as in (C)], which had themselves been pre-treated with and without anti-AREG neutralising antibody or with a control IgG. Growth curves were reproduced in three independent experiments. Bar charts derived from three independent experiments: *P < 0.05, **P < 0.01. See also Figure S6.

Since epidermal growth factor receptor (EGFR) signalling plays a key role in CRC growth and the EGF ligand Amphiregulin (AREG) is expressed by HCT116 cells (Nagathihalli et al., 2014) and known to be packaged into CRC-derived exosomes (Higginbotham et al., 2011), we tested whether AREG was present on EVs from glutamine-depleted HCT116 cells. Glutamine depletion increased levels of membrane-associated AREG (Brown et al., 1998) in cell lysates and in EVs produced under these conditions (Figure 6C, S6E). Treating EV preparations from glutamine-depleted and control cells with a neutralising antibody to AREG (Raimondo et al., 2019) blocked the growth-promoting effect of glutamine-depletion-induced EVs (Figure 6C’), suggesting that AREG-positive vesicles play a critical role in this process.

### Glutamine-Depletion-Induced Extracellular Vesicles Promote Tumour Cell Turnover *in Vivo*

To test whether the enhanced activities of EV preparations from glutamine-depleted HCT116 cells could be replicated *in vivo*, we established HCT116 xenografts in exponential growth phase and then directly injected them every three days over nine days with EVs, isolated from HCT116 cells under glutamine-replete and -depleted conditions. Xenografts were analysed 24 h following the last injection of EVs. There was no change in overall tumour growth over this short time period following exposure to either type of EV preparation (Figure S7). However, there were significant histological differences within the xenografts for different treatments. As previously reported (Sheldon et al., 2010), injecting EVs into xenografts led to increased overall vessel number (Figure 7A); however, the effect of the two EV preparations was different, with vessel lumen size significantly increased in xenografts exposed to EVs from glutamine-depleted cells. The increased number of blood vessels for both EV treatments correlated with elevated tumour cell proliferation (Figure 7B). Again, consistent with observations *in vitro*, exposure to EV preparations from glutamine-depleted cells led to the highest levels of proliferating cells in the viable regions of the tumour (Figure 7B). However, an overall increase in necrosis (Figure 7C) and in hypoxia, as indicated by expression of carbonic anhydrase (CA9) as a downstream hypoxia marker (Figure 7D; McIntyre et al., 2016), was also induced by EV treatment, and particularly by EV preparations from glutamine-depleted cells. Overall, we conclude that changes in EV production driven by glutamine depletion induce increased cell turnover *in vivo*, which may contribute to adaptive changes under these metabolic stress conditions.

**Figure 7.**
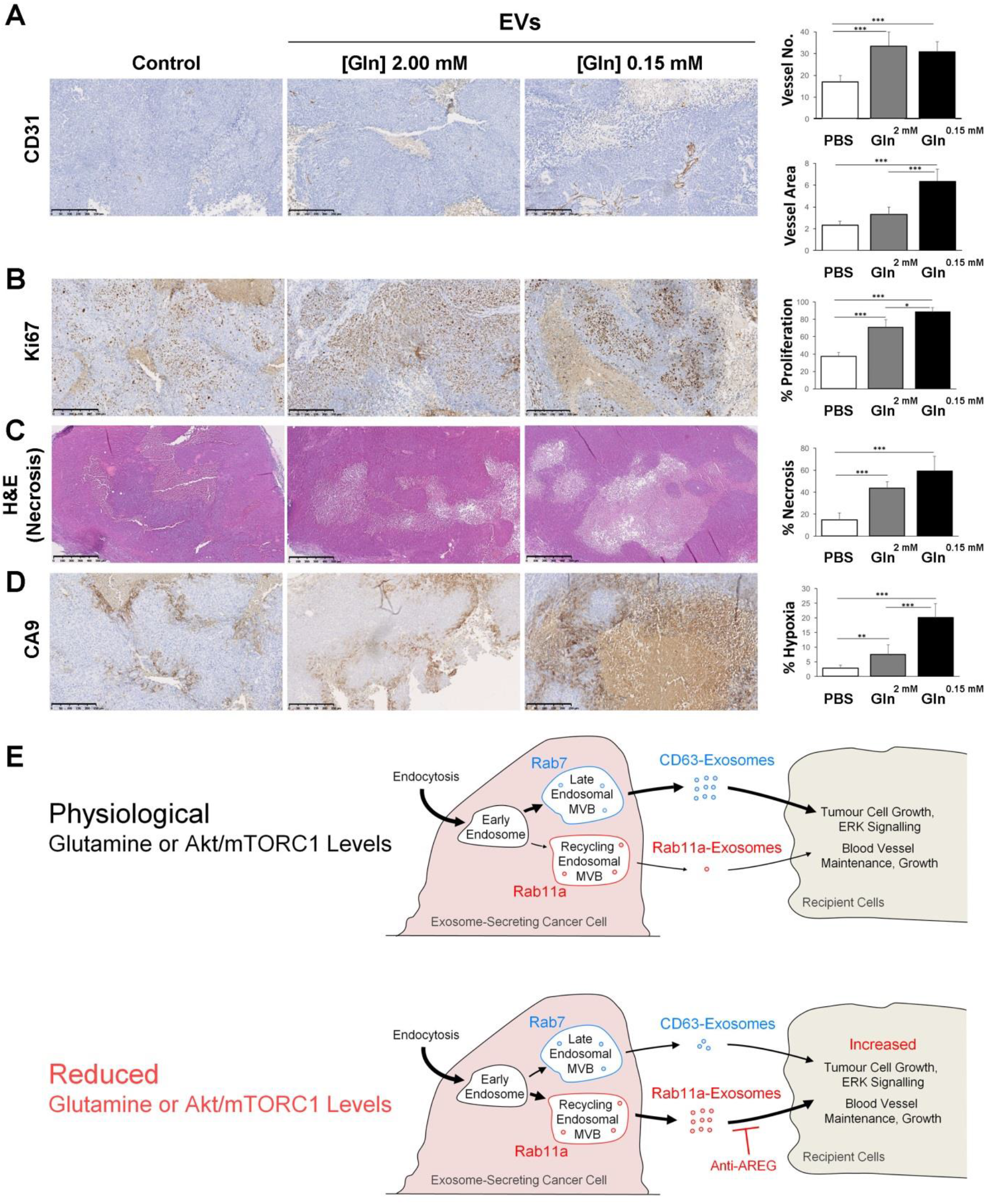
EVs from Glutamine-Depleted HCT116 Cells Increase Tumour Cell Turnover and Vessel Growth in Mouse HCT116 Xenografts. HCT116 flank tumours produced by subcutaneous injection were established for 20 days before injection at three-day intervals with vehicle (PBS), or EVs isolated by UC from glutamine-replete or glutamine-depleted HCT116 cells. Tumours were excised 24 h after last of four injections for analysis. Panels show representative immunostained histological sections of tumour tissue quantified using the Visiopharm Integrator System. (A) Sections immunostained for CD31, which labels endothelial cells and blood vessels, with blood vessel number (upper) and total area (lower) represented in bar charts. (B) Sections immunostained for Ki67, which stains proliferative cells. Proportion of tumour cells with Ki69 staining is represented in bar chart. (C) Sections stained with haematoxylin and eosin, which highlights necrotic regions. Proportion of tumour area that is necrotic is represented in bar chart. (D) Sections immunostained for CA9, which is expressed in hypoxic regions. Proportion of tumour cells with CA9 staining is represented in bar chart. (E) Schematic model showing how in cancer cells, regulation of endosomal trafficking by depletion of exogenous glutamine or reduced Akt/mTORC1 signalling can induce a switch in exosome production from the established Rab7-late endosomal mutivesicular bodies (MVBs; producing ‘CD63-exosomes’), to Rab11a-labelled recycling endosomal MVBs (generating ‘Rab11a-exosomes’). In recipient cells, the resulting vesicles can increase ERK signalling and cell growth in tumour cells, and enhance growth and stability of endothelial network formation. Thickness of arrows indicates levels of membrane flux through the two exosome-generating endosomal routes. Scale bar is 250 µm, except for (C), which is 500 µm. Data were analysed by one-way ANOVA; n = 7; * P < 0.05, ** P < 0.01, *** P < 0.001. See also Figure S7.

## DISCUSSION

Exosomes are important mediators of signalling between cells, particularly in cancer, but the mechanisms by which exosome signals might change in response to microenvironmental stress have remained largely unexplored. Here, using human CRC, cervical and prostate cancer cell lines, and supported by a fly *in vivo* exosome biogenesis model, we provide evidence that exosomes are not only formed in late endosomal MVBs, but also in Rab11/11a-positive recycling endosomal MVBs. These latter exosomes carry distinct cargos, including Rab11/11a, providing a diagnostic signature for their compartment of origin. Furthermore, they are preferentially released by cancer cells following glutamine depletion or Akt/mTORC1 inhibition, generating unique biological effects *in vitro* and *in vivo*, (shown schematically in Figure 7E). We propose that this previously undescribed exosome subtype plays a unique role in tumour adaptation to metabolic stresses.

### Rab11/11a-Positive Compartments are Novel Sites of Exosome Biogenesis

Knockdown studies in human cells and *Drosophila* have highlighted a role for Rab11 family members in exosome secretion (Savina et al., 2002; Koles et al., 2012; Beckett et al., 2013). In fact, one report previously identified Rab11a-positive MVBs in human cells (Savina et al., 2005). These experiments were, however, taken as evidence for Rab11a facilitating MVB trafficking to the cell surface and/or docking MVBs there. Our *in vivo* analysis in *Drosophila*, together with imaging studies and EV analysis in human cell lines indicates that Rab11-compartments are conserved sites of exosome biogenesis. Rab11 family members are classically associated with recycling endosomes, but are also implicated in regulating secretory traffic from the Golgi (Welz et al., 2014). Human Rab11a-positive exosomes, however, appear to be generated in recycling endosomes, because their secretion is upregulated when traffic through the recycling pathway is increased, and Rab11a-positive ILVs can form in enlarged Rab5CA-induced endosomes. High-resolution imaging in the fly system has allowed us to demonstrate that production of these vesicles is ESCRT-dependent, suggesting parallels with late endosomal exosomes in their biogenesis mechanisms.

Glutamine depletion induces a switch in EV secretion in cancer cells that includes increased release of Rab11a-exosomes. Both in *Drosophila* (Corrigan et al., 2014) and in glutamine-depleted human cells, blocking Rab11/11a activity inhibits the biological activities of secreted EVs, suggesting a key role for exosomes generated in Rab11/11a compartments. This manipulation could have other indirect effects on secretion, but at least in HCT116 cells, *Rab11a* knockdown leads to only modest or no reduction in several other exosome markers in EV preparations, and increases Cav-1 secretion. The treatment therefore appears to have relatively specific effects on Rab11a-exosome secretion and suggests these exosomes have important disease-relevant activities.

The continued release of Cav-1 following *Rab11a* knockdown in HCT116 cells suggests that its secretion under stress conditions involves another EV biogenesis pathway, either within unidentified intracellular compartments or from the cell surface. Secretion of Cav-1-containing small EVs of unknown subcellular origin has also recently been shown to be under metabolic (Crewe et al., 2018), as well as hypoxic (Kucharzewska et al., 2013) control. In the latter case, this may involve reduced constitutive endocytosis (Bourseau-Guilman et al., 2016), but a role for the Rab11a pathway was not been investigated. Distinguishing the different classes of tumour EV that are produced under metabolic stress should assist in determining what cancer-relevant functions are associated with each specific EV subtype, as we have done for Rab11a-exosomes.

Although not major constituents of exosomes, the identification of Rab11/11a and potentially Rab7 (Figure 3E) as signatures for exosome origin suggests a new approach for distinguishing different exosome subtypes in EV preparations. In flies, the selective labelling by Rab11 of only a minority of vesicles in Rab11-compartments, unlike Btl-GFP, suggests the existence of subpopulations of vesicles in a single compartment. Unlike transmembrane proteins present on the limiting membrane of secretory compartments, Rabs are thought to disengage from the lipid bilayer before or during plasma membrane fusion, as evidenced by our fly data (Figure S1B, S1C), so they are unlikely to be incorporated into microvesicles shed from the cell surface. Multiple Rabs have been reported to be present in EV preparations (Keerthikumar et al., 2016), suggesting that other exosome subtypes are yet to be identified. Of particular interest are Rab35 and Rab27 family members, which like Rab11, have been implicated in exosome release (Hsu et al., 2010; Ostrowski et al., 2010). Whether they mark additional exosome biogenesis pathways or primarily promote trafficking from other pathways remains to be determined.

### Tumour Exosome Signalling is Regulated by Metabolic Stress and Akt/mTORC1 Signalling

Our finding that several cancer cell types alter their exosome secretion and signalling following glutamine depletion highlights a novel function for the recycling endosomal exosome biogenesis pathway in a tumour’s response to its microenvironment. In contrast to cancer cells, the highly secretory SCs of the fly accessory gland release exosomes from Rab11-positive compartments containing dense-core granules in a regulated fashion under normal physiological conditions (Corrigan et al., 2014; Redhai et al., 2016). Rab11 family members have also been implicated in granule secretion from pancreatic beta cells (Sugawara et al., 2009) and *Drosophila* salivary glands (Farkaš et al., 2015), although to date, ILVs have not been reported in the much smaller compartments involved. It will be interesting to investigate whether highly secretory cells use an entirely different mechanism to control this form of secretion or whether there is an unappreciated link to metabolic stress.

Growth factor- and nutrient-dependent regulation of many cellular functions, including proliferation, apoptosis, autophagy and metabolism involves mTORC1 (Goberdhan et al., 2016). The roles of mTORC1 in secretion, particularly exosome release, have been less extensively studied. A recent report suggests that exosome secretion is increased after mTORC1 inhibition in mouse embryonic fibroblasts, but markers like Rab11a and Cav-1 were not analysed in this study (Zou et al., 2018). Our findings suggest that therapeutic blockade of PI3K/Akt or mTORC1 signalling in cancer can induce a switch in secreted exosome subtype, although in some cells, complete mTORC1 inhibition appears to strongly reduce all exosome secretion. In addition to highlighting a new role for growth factor signalling and glutamine in exosome regulation, and its potential role in adaptation mechanisms, our data indicate the importance of monitoring mTORC1 activity during EV isolation procedures.

### Exosome Signalling in Tumour Adaptation

We were initially surprised to find that the release of recycling endosomal exosomes from glutamine-depleted HCT116 cells enhances tumour cell growth, in addition to promoting blood vessel maintenance *in vitro*. Our subsequent xenograft analysis has allowed us to detect similar activities *in vivo* and develop a model that may explain these findings.

Consistent with previous reports, we observed that injecting EVs from glutamine-replete or - depleted cells increased the number of blood vessels within tumours (Sheldon et al., 2010). However, EVs from glutamine-depleted cells also preferentially increased proliferation in the tumour, which likely contributed to the elevated hypoxia and necrosis within the xenografts, despite the increased number of blood vessels. Tumour cells under nutrient and hypoxic stress are known to adapt to such metabolic stress conditions to survive and grow. For example, cancer cells can become ‘glutamine addicted’, whereby glutamine metabolism is able to sustain proliferation (Tardito et al., 2015). These cells often upregulate their growth regulatory pathways to achieve this. For instance, oncogenic KRAS drives glutamine metabolism in lung adenocarcinoma (Romero et al., 2017), while in lung squamous cell carcinoma, where mTORC1 is frequently hyper-activated, it has been observed that tumours that are highly resistant to conventional chemotherapy respond to inhibitors targeting the glutamine pathway (Lukey et al., 2018). Although we show an important role for amphiregulin in glutamine-depletion-induced growth, the HCT116 cell line already has a KRAS mutation, which may make it more likely to respond to glutamine deprivation or to the exosomes generated under such circumstances.

Release of Rab11a-exosomes under metabolic stress provides a mechanism for generating the heterogeneity in tumours that might drive adaptive changes of this kind. The secretion of growth-promoting molecules associated with Rab11a-exosomes from cells might promote their quiescence or even apoptosis. However, it could also stimulate a subpopulation of other cells that are more resistant to metabolic stress to activate ERK and other growth factor cascades, proliferate, alter their metabolism and potentially out-compete their neighbours. Our *in vitro* studies with an anti-AREG neutralising antibody suggest that AREG-EGFR signalling may be involved in this process, although we cannot eliminate the possibility that binding of this antibody to AREG-containing exosomes indirectly interferes with other exosome functions.

Since glutamine is now recognised as a key metabolite in several cancers (Zhang et al., 2017), glutamine depletion is likely to be a common event in growing tumours, which would trigger a switch to Rab11a-exosome secretion. Reducing growth factor and associated mTORC1 signalling could have similar effects. Therefore, blocking this form of Rab11a-exosome communication may have therapeutic value in suppressing adaptation and/or resistance mechanisms, a concept that needs to be explored with further *in vivo* experiments. In addition, the use of exosomes as biomarkers for cancer and other diseases has considerable clinical potential (Barile and Vassalli, 2017, Shah et al., 2018). Our data suggest that the tumour exosome profile can indicate the response of cancer cells to microenvironmental stresses, anti-angiogenic drugs and inhibitors of Akt/mTORC1 signalling. A better understanding of metabolically regulated exosome secretion should therefore lead to exosome profiling being applied with increasing precision in future diagnosis and in prognostic applications involving such treatments.

## MATERIALS AND METHODS

### *Drosophila* Stocks and Genetics

The following fly stocks, acquired from the Bloomington *Drosophila* Stock Centre, unless otherwise stated, were used: *UAS-CD63-GFP* (Panáková et al., 2005; a gift from S. Eaton, Max Planck Institute of Molecular Cell Biology and Genetics, Dresden, Germany), *dsx-GAL4* (Rideout et al., 2010; a gift from S. Goodwin, University of Oxford, UK), *tubulin-GAL80^ts^*; *UAS-Btl-GFP* (Sato and Kornberg, 2002), *YFP^MYC^-Rab11* and *YFP^MYC^-Rab7* (Dunst et al., 2015, a gift from F. Karch and E. Prince, Department of Genetics and Evolution, University of Geneva, Switzerland), *UAS-shrb-RNAi* (RNAi [1] - v106823: Dietzl et al., 2007, Matusek et al., 2015; Vienna *Drosophila* Resource Centre (VDRC); *UAS-Vps28-RNAi* (v31894; Dietzl et al., 2007, Matusek et al., 2015; VDRC), *btl-Gal4* (*btl^NP6593^*; Hayashi et al., 2002; Kyoto *Drosophila* Stock Centre); *UAS-GFP^NLS^*; *UAS-YFP-Rab11* (Zhang et al., 2007), *UAS-shrb-GFP* (Sweeney et al., 2006); and *w^1118^* (a gift from L. Partridge, UCL, UK).

Flies were reared at 18°C in vials containing standard cornmeal agar medium containing, per litre, 12.5 g agar, 75 g cornmeal, 93 g glucose, 31.5 g inactivated yeast, 8.6 g potassium sodium tartrate, 0.7 g calcium, and 2.5 g Nipagen (dissolved in 12 ml ethanol). They were transferred onto fresh food every three to four days. No additional dried yeast was added to the vials.

Temperature-controlled, SC-specific expression of *UAS-CD63-GFP*, *UAS-Btl-GFP* and *UAS-Rab11-YFP* was achieved by combining these transgenes with the specific driver, *dsx-GAL4* and the temperature-sensitive, ubiquitously expressed repressor *tubulin-GAL80^ts^*. Newly eclosed virgin adult males were transferred to 28.5°C to induce post-developmental SC-specific expression. For knockdown experiments, the same strategy was employed, but in the presence of a UAS-RNAi transgene. Six-day-old adult virgin males were used throughout this study to ensure that age- and mating-dependent effects on SC biology were mitigated (Leiblich et al., 2012; Redhai et al., 2016).

To produce a pulse of Shrb-GFP expression, newly eclosed virgin males were kept at 25°C (permissive for GAL80^ts^ activity) for six days, before being transferred to 28.5°C for a 4 h ‘pulse’ of transgene expression under *dsx-GAL4* control, then returned to 25°C for a further 4 h before imaging.

### Preparation of Accessory Glands for Live Imaging

Adult male flies were anaesthetised using CO_2_. The anaesthetised flies were submerged in ice-cold PBS (Gibco^®^) during the micro-dissection procedure. The whole male reproductive tract was carefully pulled out of the body cavity. The testes were then removed by scission close to the seminal vesicles, as they often folded over the accessory glands, obscuring imaging. Care was taken to keep the seminal vesicles and vas deferens intact in order to prevent damage to the papilla of the anterior ejaculatory duct. The accessory gland epithelium can be stressed by damage to this valve-like papilla, through which the spermatozoa and accessory gland contents are transferred during mating. Finally, the remaining reproductive tissues, including the accessory glands were isolated by separation of the ejaculatory bulb from the external genitalia, fat tissues and gut.

The isolated accessory glands were kept in PBS on ice until sufficient numbers had been obtained (typically for approximately 15 min). They were then stained with ice-cold 500 nM LysoTracker^®^ Red DND-99 (Invitrogen^®^) in 1 × PBS for 5 min. Finally, the glands were washed in ice-cold PBS for 1 min before being mounted onto High Precision microscope cover glasses (thickness No. 1.5H, Marienfeld-Superior). A custom-built holder secured the cover glasses in place during imaging with both the super-resolution 3D-SIM and wide-field systems. To avoid dehydration and hypoxia, the glands were carefully maintained in a small drop of PBS, surrounded by 10S Voltalef^®^ (VWR Chemicals), an oxygen-diffusible hydrocarbon oil (Parton et al., 2010), and kept stably in place by the application of a small cover glass (VWR).

## Imaging

### Confocal Imaging

Confocal images of cultured cells and accessory glands were acquired either by using a Zeiss LSM 510 Meta [Axioplan 2] or LSM 880 laser scanning confocal microscope equipped with a 63x, NA 1.4 Plan APO oil DIC objective (Carl Zeiss). RI 1.514 objective immersion oil (Carl Zeiss) was employed.

### Super-Resolution 3D-SIM Microscopy

For super-resolution 3D-SIM, fixed human cells or freshly prepared living *Drosophila* accessory glands were imaged on a DeltaVision OMX V3 system (GE Healthcare Life Sciences) equipped with a 60x, NA 1.42 Plan Apo oil objective (Olympus), sCMOS cameras (PCO), and 405, 488, 593 and 633 nm lasers. Objective immersion oil of RI 1.516 (Cargille Labs) was used to produce optimal reconstruction for a few μm along the Z-axis of the sample.

The microscope was kept at approximately 20°C throughout experiments. Images were acquired in sequential imaging mode, with Z-stacks, typically spanning 8-12 µm, being acquired with 15 images per plane (five phases, three angles) and a z-distance of 0.125 µm. The raw data were computationally reconstructed in the softWoRx 6.0 software package (GE Healthcare Life Sciences), using a Weiner filter of 0.002 and wavelength-specific optical transfer functions (OTFs) to generate super-resolution images with two-fold improved resolution in all three axes. The ImageJ2 plugin ‘SIMcheck’ (Ball et al., 2015) was used to analyse the quality of the reconstructed images. As the OMX system uses separate cameras for each colour, the originally produced images required careful re-alignment. To make the appropriate adjustments, reference Z-stacks of 200 nm TetraSpeck Microspheres (Invitrogen^®^, T7280) were aligned using custom automated software (OMX Editor) that corrected for X, Y and Z shifts, magnification and rotation differences between channels. The resulting transformations were then applied to align the images of the test samples.

### Wide-Field Microscopy

For wide-field imaging, living SCs were imaged using a DeltaVision Elite wide-field fluorescence deconvolution microscope (GE Healthcare Life Sciences) equipped with both a 100x, NA 1.4 UPlanSApo oil objective (Olympus) and a 60×, NA 1.42 UPlanSApo oil objective (Olympus), as well as a CoolSNAP HQ2 CCD camera (Photometrics^®^) and a seven colour illumination system (SPECTRA light engine^®^, Lumencor). A 1.6x auxiliary magnification lens and objective immersion oil with RI 1.514 (Cargille Labs) were used throughout. The microscope room was temperature-controlled at ∼18°C. The images acquired were typically Z-stacks spanning 8-12 µm depth with a z-distance of 0.2 µm. Images were subsequently deconvolved using the Resolve 3D-constrained iterative deconvolution algorithm within the softWoRx 5.5 software package (GE Healthcare Life Sciences).

### Electron Microscopy

To image isolated EVs, 10 µl of a purified EV suspension were incubated on a glow discharged 300 mesh copper grid coated with a carbon film for 2 min, blotted with filter paper and stained for 10 s with 2% uranyl acetate. The grid was then blotted and air-dried. Grids were imaged in a Tecnai FEI T12 transmission electron microscope (TEM) operated at 120 kV with a Gatan Oneview digital camera.

For SC studies, a High Pressure Freezing (HPF) and freeze substitution method was employed. Accessory glands from six-day-old virgin *w^1118^* males were carefully dissected in ice-cold PBS under a dissection microscope. The glands were then removed from the PBS and coated in yeast paste (dry baker’s yeast mixed with 10% methanol), which acts as a cryo-protectant. Four or five glands were transferred using forceps into the well of a membrane carrier (1.5 mm diameter, 100 μm deep, Leica). The carriers were then loaded into bayonet pods (Leica) and transferred using a manual loading device to the high pressure freezer (EM PACT2, Leica). Samples were frozen immediately at 2000 bar.

The carriers were carefully moved into pre-cooled 2 ml screw top cryotubes containing pre-cooled freeze substitution medium (0.1% uranyl acetate and 1% osmium tetroxide in dry acetone) within the sample chamber of an automatic freeze substitution unit (EM AFS2, Leica) held at −130°C. Over the following 40 h, the samples were gradually brought up to room temperature to avoid formation of ice crystals.

Following freeze substitution, the samples were then moved using a plastic Pasteur pipette into fresh 2 ml tubes of 100% acetone. Samples were then gradually infiltrated with Agar 100 Hard resin (Agar Scientific) at room temperature over several days. Samples were first incubated with a 3:1 mix of 100% dry acetone:resin (without accelerator) for 2 h, then a 1:1 mix for 3 h, then a 1:3 mix for 2 h. The samples were then left overnight in 100% resin (with accelerator from now on), which was replaced with fresh 100% resin four times over 32 h. The glands were carefully transferred to and positioned in embedding capsules, and the resin polymerised at 60°C in an oven over 40 h. Ultra-thin (90 nm) sections were taken with a Diatome diamond knife on a Leica UC7 ultramicrotome and transferred to 200 mesh copper grids, which were then post-stained for 5 min in uranyl acetate, washed 8 times in water and then stained for 5 min in lead citrate. The grids were washed seven times in water, and allowed to dry. Samples were imaged as described for EV preparations above.

### *Drosophila* Exosome Secretion Assay

Control virgin six-day-old SC>CD63-GFP- or SC>Btl-GFP-expressing males and virgin CD63-GFP-/Btl-GFP males also expressing an RNAi against ESCRT components *shrb* or *Vps28* in SCs were dissected in 4% paraformaldehyde (Sigma-Aldrich) dissolved in PBS. The glands were left in 4% paraformaldehyde for 20 minutes to preserve the luminal contents before being washed in PBS for 5 min. Glands were then mounted onto SuperFrost^®^ Plus glass slides (VWR), removing excess liquid using a tissue, and finally immersed in a drop of Vectashield^®^ with DAPI (Vector Laboratories) for imaging by confocal microscopy.

Exosome secretion was measured using a modification of a previously described approach (Corrigan et al., 2014), in which multiple areas of luminal fluorescent puncta (exosomes or exosome aggregates) were sampled within the central third of each gland. Identical microscope settings and equipment were used throughout. At each sampling location, three different Z-planes were analysed, spaced by 5 μm for CD63-GFP, and ten different z-planes were analysed, spaced by 1 μm for Btl-GFP, with the first image in the Z-stack located exactly 5 μm apical to the main cell nuclei for both markers. This helped to ensure similar fluorescence levels between glands of the same genotype.

The automated analysis of exosome secretions by SCs was performed using ImageJ2 (Schindelin et al., 2015), distributed by Fiji. The raw data from each lumenal area were thresholded using the Kapur-Sahoo-Wong (Maximum Entropy) method (Kapur et al., 1985) to determine the outlines of fluorescent particles. The ‘Analyse Particles’ function was then used to determine the total number of fluorescent puncta.

### Analysis of ILV Content in Large Non-Acidic Compartments

Living SCs from six-day-old SC>Btl-GFP-, SC>CD63-GFP-, or SC>YFP-Rab11-expressing flies (with or without RNAi expression) were imaged using identical settings by wide-field microscopy. The total number of fluorescently labelled non-acidic compartments and the total number of these compartments that contained fluorescently labelled puncta was counted in each cell, using a full Z-stack of the epithelium. Three individual SCs were analysed from each of 13 glands.

### SC Compartment Analysis

SC compartment numbers were manually analysed in ImageJ2 using the ‘Cell Counter’ plugin. Complete Z-stacks of individual cells were scored for the number of large non-acidic compartments (LysoTracker Red^®^-negative) and the number of large acidic late endosomal and lysosomal (LEL) compartments (LysoTracker Red^®^-positive). The diameter of the largest LEL was measured manually at the Z-plane where the diameter was at its greatest using the line tool (see Corrigan et al., 2014; Redhai et al., 2016). Both fluorescently labelled and non-transgenic *w^1118^* flies were analysed using a combination of fluorescence and DIC microscopy on the wide-field microscope. Three individual SCs per gland from each of ten glands were used to calculate the mean compartment number in each genotype.

### Measurement of ILV Diameter

Super-resolution 3D-SIM images of living SCs expressing CD63-GFP were used to determine the diameter of individual ILVs within non-acidic compartments. Careful manual measurements were taken of the diameters of ILVs within individual compartments using the line tool in ImageJ2. Many ILVs were smaller than the resolution limits of super-resolution 3D-SIM, and were visualised as individual bright puncta (<100 nm diameter). The three largest compartments per SC were analysed by this method and the mean calculated from three separate SCs in each of three glands.

### Cell Culture

HCT116, HeLa and LNCaP cells were purchased from ATCC. Cells were grown in McCoy’s 5A modified medium (Life Technologies; HCT116), DMEM medium (Life Technologies; HeLa) or RPMI-1640 medium (Life Technologies; LNCaP), supplemented with 10% heat-inactivated foetal calf serum (FBS), 100 U/ml Penicillin-Streptomycin (Life Technologies), and incubated in 5% CO_2_ at 37°C prior to EV collection.

### Extracellular Vesicle Isolation

During EV isolation, HCT116 and HeLa cells were maintained in serum-free basal medium (DMEM/F12) supplemented with 1% ITS (Insulin-Transferrin-Selenium; #41400045 Life Technologies) (Tauro et al., 2012) for 24 h. LNCaP cells were cultured in RPMI-1640 supplemented with 1% ITS for 24 h. For glutamine depletion experiments, cells were grown for 24 h in DMEM/F12 medium without L-glutamine (#21331046; Life Technologies) supplemented with 1% ITS and 2.00 mM or low glutamine (0.15 mM for HCT116; 0.02 mM for HeLa and LNCaP) (Life Technologies). Torin 1 was added at 100 nM (HCT116) or 120 nM (LNCaP), rapamycin at 10 nM, and AZD5363 at 3 µM. Generally, about 8-9 x 10^6^ HCT116 cells, 4 x 10^6^ HeLa cells, or 12 x 10^6^ LNCaP cells were seeded per 15 cm cell culture plate (typically 10-15 plates used per condition) and allowed to settle for 16-24 h in complete medium before a 24 h EV collection. Cells were typically 80-90% confluent at the end of the collection period. The culture medium was pre-cleared by centrifuging at 500g for 10 min at 4°C and 2,000g for another 10 min at 4°C to remove cells, debris and large vesicles. The supernatant was filtered using 0.22 μm filters (Milex^®^).

For EV isolation by differential ultracentrifugation (UC), large volumes of the EV-containing filtrate were concentrated using a tangential flow filter (TFF) set-up with a 100 kDa membrane (Vivaflow 50R, Sartorius) using a 230V pump (Masterflex). The EV suspension was typically centrifuged at 100,000g - 108,000g for 70 - 120 min in a Beckman 55.2 Ti or 70 Ti rotor at 4°C (Beckman Coulter). For activity assays and some experiments involving western analysis, the EV pellet was resuspended in 25 ml PBS and the ultra-centrifugation step repeated to minimize soluble protein contaminants. Finally, the EV pellet was resuspended in 70-100 µl PBS for subsequent experiments. For activity, Proteinase K and most NTA assays, EVs were used within 24 h, but for other analyses, including EVs used for xenograft injections, they were stored at −80°C prior to use on the day of the injection experiment.

To isolate EVs by size-exclusion chromatography, EV-containing filtrate from a 2000g centrifugation was concentrated using a 100 kDa TFF filter to a volume of approximately 30 ml and then further concentrated in 100 kDa Amicon filters by sequential centrifugation at 4000g for 10 min at 4°C to a final volume of 1 ml. This was injected into a size-exclusion column (column size 24 cm x 1 cm containing Sepharose 4B, 84 nm pore size) set up in an AKTA start system (GE Healthcare Life Science) and eluted with PBS, collecting 30 x 1 ml fractions. Fractions corresponding to the initial “EV peak” (fractions two to five for HCT116 and fractions two to seven for LNCaP cells, which appeared before the large protein peak) were pooled in 100 kDa Amicon tubes to a final volume of approximately 100 µl for analysis.

Initially, EV preparations for western and functional analysis were assessed after normalisation based on the mass of protein in EV-secreting cells and on EV number, determined by NTA. To normalise EV preparations to the mass of protein in EV-secreting cells, cells were washed in PBS twice and then lysed on ice in cold RIPA buffer supplemented with protease and phosphatase inhibitors (Sigma). Cell debris was pelleted by centrifugation at 13,000g for 10 min at 4°C. Protein concentration in the supernatant was measured by bicinchoninic acid (BCA) assay (Pierce, Thermo Scientific). Although glutamine depletion did not significantly affect EV number and the normalisation method did not affect the conclusions from our analysis (Figures 4B, S4E, S5H), after some drug and knockdown treatments, EV production was reduced by up to 50% (Figure S5H). We elected to analyse EVs based on EV-secreting cell mass to ensure that any changes in protein levels did not result from testing EVs produced by altered number of cells.

EVs were isolated by ultrafiltration using a modification of the protocol described by Cheruvanky et al., (2007). The 0.22 μm filtrate, containing EVs (typically 40 ml volume), was concentrated in a 100 kDa Amicon Ultra-15 (Millipore^®^) filter by sequential centrifugation of approximately 15 ml aliquots at 4000g for 10 min at 4°C. The EVs were washed in PBS twice, by repeating the centrifugation step, to remove soluble proteins. The EV suspension in approximately 140-170 µl PBS was removed from the filter by pipette. This method gave more consistent recovery of EV proteins for western analysis, although results were always confirmed using EVs isolated by UC. The method was not suitable for activity or NTA assays, but was useful for some preliminary analysis and for processing multiple samples simultaneously.

To test whether EV proteins were located inside vesicles, ultracentrifugation-isolated EV samples were resuspended in Dulbecco’s PBS (DPBS; Gibco^®^, #14040-083), then incubated with or without proteinase K at 25 µg/ml (Roche, #03115828001), and with or without 0.1% Triton-X (Sigma, #T8787) at 37°C for 30 min. Proteinase K digestion was terminated using 5 mM PMSF (Cell Signalling Technology, #8553S) and the samples subjected to Western blot analysis.

### Nanoparticle Tracker Analysis (NanoSight**^®^**)

The NS500 NanoSight^®^ was used. Thirty sec videos were captured between three and five times per EV sample at known dilution (normalised to protein mass of secreting cells). Particle concentrations were measured within the linear range of the NS500 between about 2-10 x 10^8^ particles per ml. Particle movement was analysed by NTA software 2.3 (NanoSight Ltd.) to obtain particle size and concentration.

### Genetic Manipulation of Cultured Cells

For shRNA knockdown, the following conditions were used. For *Rab11a*, cells were cultured for two days following transfection either with *Rab11a* shRNA lentivirus (AAGAGCACCATTGGAGTAGAG; virus generated using pHR-U6 vector) or new shRNA lentivirus (GTTGTCCTTATTGGAGATTC; TRCN0000381243; Sigma) before EV collection. For *raptor*, cells were cultured for three days following transfection with *raptor* shRNA lentivirus (Addgene #1858; Sarbassov et al., 2005) before EV collection. For *Rab7a*, cells were cultured for three days following transfection with *Rab7* shRNA lentivirus (GGCTAGTCACAATGCAGATAT; in pHR-U6 vector) before EV collection. For NT shRNA, the shRNA sequence was CAACAAGATGAAGAGCACCAA (SHC202, Sigma).

Transient transfections of fusion protein constructs were carried out when the cells reached 80% confluency with the transfection reagent Lipofectamine^®^ 2000 in serum-reduced OptiMEM^®^ medium (Gibco^®^, Life Technologies), and following manufacturers’ instructions to obtain maximum transfection rates. Constructs were GFP-Rab5CA (Q79L) (Addgene #35140) and GFP-Rab11 WT (Addgene #12674).

### Western Analysis

Both cell lysates and EV preparations were electrophoretically separated using 10% mini-PROTEAN^®^ precast gels (BioRad). EV preparations were lysed in RIPA or 1X sample buffer. EV loading was normalised to cell lysate protein levels to take account of changes in cell number in different conditions. Protein preparations were ultimately dissolved in either reduced (containing 5% β-mercaptoethanol) or non-reduced (for CD63 and CD81 detection) sample buffer and were heated to 90-100°C for 10 min before loading with a pre-stained protein ladder (Bio-Rad). Proteins were wet-transferred to Polyvinylidene difluoride (PVDF) membranes at 100 V for 1 h using a Mini Trans-Blot Cell (Bio-Rad). The membranes were then blocked with either 5% milk (optimal for CD63 detection) or 5% BSA in TBS buffer with Tween-20 (TBST) for 30 min and probed overnight at 4°C with primary antibody diluted in blocking buffer. After 3 x 10 min washing steps with TBST, the membranes were probed with the relevant secondary antibodies for 1 h at 22°C. Following 3 x 10 min wash steps, the signals were detected using the enhanced chemiluminescent detection reagent (Clarity^®^, BioRad) and the Touch Imaging System (BioRad). Relative band intensities were quantified by ChemiDoc^®^ software (Bio-Rad) or ImageJ. Signals were normalised to tubulin (cell lysates), or CD81 (HCT116, HeLa and LNCaP EVs). Note, as shown in Figure 4B and S4E, that if EV protein signals are normalised to cell lysate levels, CD81, Syn-1 or Tsg101, relative levels of Rab11a (and usually Cav-1) are increased after glutamine depletion or partial blockade of mTORC1 signalling, while CD63 is typically reduced.

Antibody suppliers, catalogue numbers and concentrations used were: rabbit anti-4E-BP1 (Cell Signaling Technology #9644, rabbit anti-p-4E-BP1-Ser65 (Cell Signaling Technology #9456, 1:1000), rabbit anti-S6 (Cell Signaling Technology #2217, 1:4000), rabbit anti-p-S6-Ser240/244 S6 (Cell Signalling Technology #5364, 1:4000), rabbit anti-Raptor (Cell Signalling Technology #2280, 1:1000), rabbit anti-Caveolin-1 (Cell Signaling Technology #3238, 1:500), goat anti-AREG (R&D Systems #AF262, 1:200), mouse anti-Tubulin (Sigma #T8328, 1:4000), rabbit anti-GAPDH (Cell Signaling Technology #2118, 1:2000), mouse anti-CD81 (Santa Cruz #23962, 1:500), mouse anti-CD63 (BD Biosciences # 556019, 1:500), rabbit anti-Syntenin-1 antibody (Abcam ab133267, 1:500), rabbit anti-Tsg101 (Abcam ab125011, 1:500), mouse anti-Rab11 (BD Biosciences #610657, 1:500), rabbit anti-p44/42 MAPK (ERK; Cell Signaling Technology #4695, 1:1000), rabbit anti-p-p44/42 MAPK (Cell Signaling Technology #4370, 1:1000), rabbit anti-PRAS40 (Cell Signaling Technology #2691, 1:1000), rabbit anti-p-PRAS40-Thr246 (Cell Signaling Technology #2997, 1:1000), rabbit anti-Rab7 (Cell Signaling Technology #9367, 1:1000), rabbit anti-Calnexin (Abcam #ab213243, 1:1000), mouse anti-GM130 (Novus Biologicals H00002801-B01P), anti-mouse IgG (H+L) HRP conjugate (Promega #W4021, 1:10000), anti-rabbit IgG (H+L) HRP conjugate (Promega #W4011, 1:10000), anti-goat IgG (H+L) HRP conjugate (R&D Systems #HAF109, 1:100).

### Immmunostaining of Cultured Human Cells

For immunofluorescence studies, cultured cells on coverslips were fixed in 4% paraformaldehyde for 15 min at ∼23°C, washed 3 x 10 min in PBS, and permeabilized with 0.05% TritonX-100/PBS for 30 sec. After three PBS washes, cells were incubated with the primary antibody prepared in 5% donkey serum/PBS in a humid chamber at 4°C overnight. The coverslips were washed 4 x 10 min in PBS, then incubated with the appropriate secondary antibodies diluted in 5% donkey serum/PBS for 1 h at ∼23°C. Following 4 x 10 min PBS washes, the coverslip was mounted on a glass slide with DAPI-containing mounting medium (VectorShield; Vector Laboratories) and sealed with nail polish. Antibody concentrations were: mouse anti-Rab11 (BD Biosciences #610657, 1:50), mouse anti-CD63 (BD Biosciences #556019, 1:100), rabbit anti-Rab7 (Cell Signaling Technology #9367, 1:50), GFP-Booster ATTO 488 (Chromotek gba488-100, 1:500), donkey anti-rabbit IgG (H+L) (DyLight^®^ 550) (Abcam ab96892, 1:500), donkey anti-mouse IgG (H+L) (Alexa Fluor^®^ 488 AffiniPure; Jackson ImmunoResearch #715-545-151, 1:500).

Cell viability was measured by Trypan Blue staining. HCT116 cells were trypsinised and then resuspended in McCoy’s A medium (Gibco^®^, #26600080). The cell suspension was mixed with Trypan Blue solution (Sigma, #T8154) at 1:1 ratio and the number of blue-staining (non-viable) cells and total cell number were assessed using a haemocytometer.

### Growth, Proliferation, Apoptosis and Neutralising Antibody Assays

For growth assays, 2 x 10^3^ HCT116 cells per well were seeded in a 96-well plate in 100 µl of medium, following pre-treatment with freshly prepared EVs for 30 min in PBS at 37°C (up to 10^4^ EVs [isolated by UC or SEC] per cell for controls, which is roughly the number of EVs secreted by each HCT116 cell in 24 h). The cells were maintained in McCoy’s 5A medium in reduced serum (1% FBS) conditions. Growth was followed over a period of five days by live image acquisition using the IncuCyte ZOOM^®^ analysis system (10X magnification; Essen Bioscience) to automatically detect cell edges and to generate a confluence mask for cell coverage area calculation. Each biological replicate is represented as mean of at least eight technical replicates.

For ERK inhibition experiments, SCH772984 (S7101; SelleckChem) was added to the cell suspension along with the exosomes to a final concentration of 1 µM. The medium was changed 24 h after cell plating.

To assess cell proliferation, cells were fixed with methanol and stained with DAPI. The number of nuclei/field for three fields per well was counted manually from fluorescence microscope images. To measure levels of apoptosis, Caspase-3 and −7 activities were assayed using 1 μM of Cell Player^®^ Caspase-3/7 reagent (Essen Bioscience), monitored with the IncuCyte ZOOM^®^ system.

AREG neutralising antibody experiments were performed in a 96-well plate using 2 x 10^3^ HCT116 cells per well treated with 4000 SEC-isolated EVs per recipient cell, using goat anti-AREG (R&D Systems #AF262, 1:200), and goat IgG (R&D Systems, AB-108-C) as a control antibody treatment. Prior to mixing with recipient cells, EVs were pre-incubated with 4ug/ml anti-AREG or control antibody in McCoy’s 5A medium with 1% FBS for 2 h at 37°C. The mixture was then added to the recipient cells (also in reduced serum media) and allowed to incubate for a further 30 min before plating. Growth was monitored using the IncuCyte ZOOM^®^ analysis system in the same fashion as standard growth assays described above.

All assays were repeated on at least three independent occasions.

### Tubulation Assay with HUVEC Cells

HUVEC cells were suspended in LONZA medium (Endothelial Basal Medium-2 - EBM2) containing freshly prepared exosomes (10^4^ UC-isolated EVs per cell). After 10 min incubation, 6 x 10^3^ cells were seeded onto a solidified thin layer of BD phenol red-free Matrigel (BD Bioscience) in a 96-well plate and incubated at 37°C. Images of vessel growth were captured at 3 h intervals for 18 h by live image acquisition using the IncuCyte ZOOM^®^ microscope (10x lens). Quantification of tubule length was performed using ImageJ. Each biological replicate is represented as mean of at least eight technical replicates.

### Assessment of EV-Induced ERK Activation

To assess ERK activation upon EV treatment, HCT116 cells were plated in six-well plates. After 24 h the cells were serum-starved for an additional 18 h. At this point, freshly prepared exosomes (10^4^ UC-isolated EVs per cell) were added and the mixture incubated for 30 min at room temperature. Cell lysates were generated and analysed by western blot.

### Cellular Uptake Assays

For confocal analysis of Tfn uptake, 5 x 10^4^ HCT116 cells were plated on a coverslip in McCoy’s 5A medium supplemented with 10% FBS and cultured overnight, followed by a 24 h incubation in DMEM/F-12, 1% ITS, with 2.00 mM or 0.00 mM glutamine at 37°C. The cells were then washed using cold PBS and cultured in DMEM/F-12, 1% ITS, containing transferrin-Alexa Fluor^®^ 488 (ThermoFisher Scientific #T13342, 1:200) with the appropriate glutamine concentration at 4°C for 30 min, followed by a 30 min incubation at 37°C. The cells were fixed for confocal imaging. For the anti-CD63 antibody uptake experiments, cells washed with cold PBS were cultured in DMEM/F-12, 1% ITS, containing mouse anti-CD63 antibodies (BD Biosciences # 556019, 1:50) and the appropriate glutamine concentration at 4°C for 30 min. The cells were then washed with PBS, prior to 30 min incubation at 37°C and a subsequent 30 min chase in DMEM/F-12, 1% ITS with the appropriate glutamine concentration. The cells were fixed, incubated with secondary antibody for the anti-CD63 uptake experiments, then imaged by confocal microscopy.

For Western analysis, 2 x 10^5^ HCT116 cells were plated in each well of a six-well plate in serum-supplemented McCoy’s 5A medium. After 20 h, the culture medium was replaced by DMEM/F-12, 1% ITS containing 2.00 mM or 0.00 mM glutamine. After 24 h, the cells were shifted to 4°C for 10 min to stop intracellular vesicular trafficking and then incubated in DMEM/F-12, 1% ITS, containing mouse anti-CD63 antibodies (1:50) and 50 µg/ml transferrin-biotin (Sigma #T3915) with appropriate glutamine concentration (2.00 or 0.00 mM) at 4°C for 30 min. After washing with cold PBS, the cells were incubated in DMEM/F-12, 1% ITS, with appropriate glutamine concentration at 37°C for 30 min and then lysed using RIPA buffer.

### Generation and Histological Analysis of HCT116 Tumours in Xenograft Mouse Models

HCT116 xenografts were established by subcutaneous injection of 2.5 x 10^6^ cells into the flank of seven female CD1 nude mice per group. When tumours had grown to an average group tumour volume of 100 mm^3^, then mice were treated with four intratumoral injections (1×10^8^ EVs/100μl exosome suspension in PBS or PBS vehicle) at three day intervals, as previously described (Sheldon et al., 2010). Tumour growth was measured three times a week using callipers and volume calculated using the formula 1/6 π x length x width x height. Twenty-four hours after the last injection, tumours were excised and half of the tumour fixed in 4% formalin overnight at 4°C before being processed and embedded in paraffin wax and sectioned for histological evaluation.

A standard haematoxylin and eosin protocol was followed to assess the morphology and the amount of necrosis in xenografts. For immunohistochemistry staining, slides were dewaxed and antigen retrieval performed at pH 6. Slides were stained using the FLEX staining kit (Agilent). Endogenous peroxidase activity was blocked before slides were stained with antibodies diluted 1:100, namely rabbit anti-Ki67 (M7240; Dako, Glostrup, Denmark), rabbit anti-CA9 (M75; BD Biosciences), and mouse anti-CD31 (JC70, Dako) for 1 h at 22°C. Slides were incubated with the appropriate anti-rabbit/anti-mouse HRP-coupled secondary antibody (Dako) for 30 min at room temperature and washed in Flex buffer and 3,3′-Diaminobenzidine (Flex-DAB; Dako) was applied to the sections for 10 min. The slides were counterstained by immersing in Flex-hematoxylin solution (Dako) for 5 min, washed and air-dried before mounting with mounting medium (Sigma). Secondary-only control staining was done routinely, but was consistently negative.

Expression of CA9 and Ki67, as well as vessel number and size, and viable/necrotic areas were quantified on whole sections quantitatively by using the Visiopharm Integrator System. The HDAB-DAB colour deconvolution band was used to detect positively stained cells. Threshold classification was used to identify necrotic and living regions, and then identify number of positive-staining cells within these regions. Threshold levels were checked against control xenograft staining before being set and the xenografts from all groups were then analysed in the same way. Vessel number and size were determined as part of a post-processing step, where areas surrounded by CD31-positive endothelial cells were automatically filled in and the areas generated counted and quantified.

### Statistical Analysis

Before parametric tests were employed, the Shapiro-Wilk Test was used to confirm normality of the datasets. For larger datasets, Levene’s Test was used to evaluate equality of variances. For Western blots, relative signal intensities were analysed using the Kruskal-Wallis test. For the growth and tubulation assays, data were analysed by two-way ANOVA. For SC analysis, a one-Way ANOVA Test was employed for normally distributed data, with post-hoc Dunnett’s Two Tailed T Tests used to directly compare individual control and experimental datasets. Non-parametric data were analysed using a Kruskal-Wallis test. We used power calculations based on previous experiments to ensure the appropriate numbers of animals were used. In order to provide a statistical power of at least 80% to detect a two-fold difference in the mean tumour volume between groups, with tumour volume variation of up to 50% within groups, and a statistical difference level of α = 0.05, a sample size of seven mice per group is required.

## ACKNOWLEDGEMENTS

We thank M. W. S. Perera for initiating studies leading to this work, and C. P. Alves for his contributions to the work presented. We are grateful to I. Dobbie, and to A. Pielach for technical expertise and support with microscopy, which was undertaken in the Wellcome Trust-funded MICRON Oxford Advanced Bioimaging Unit and Dunn School EM Facility. We thank S. Eaton, S. Goodwin, E. Prince and F. Karch, as well as the Bloomington, Vienna and Kyoto Stock Centres for *Drosophila* stocks; J. Whitburn for cells; the Developmental Studies Hybridoma Bank (Iowa) for antibodies. We are particularly grateful to K. E. Carr for comments on the manuscript. We acknowledge the support of Cancer Research UK (C19591/A19076, C602/A18974), the Cancer Research UK Oxford Centre Development Fund (C38302/A12278), the BBSRC (BB/K017462/1, BB/L007096/1, BB/N016300/1, BB/R004862/1), the Breast Cancer Research Foundation (ANR 00162), the Wellcome Trust (Strategic Awards #091911, #107457 and 102347/Z/13/Z), the National Institute for Health Research (NIHR) Oxford Biomedical Research Centre (BRC) and the John Fell Fund, Oxford (141/063).

## AUTHOR CONTRIBUTIONS

Experimental design conceptualization, S-J.F., B.K., P.M., E.B., J.D.M., K.M., C.Z., H.S., N.K.A., E.J., M.E., M.I.S., C.C.M., S.M.W., C.C., F.C.H., J.F.M., A.L.H., C.W., D.C.I.G.; experimental work, S-J.F, B.K., P.M., E.B., J.D.M., K.M., C.Z., E.J., J.F.M; data analysis, S-J.F., B.K., P.M., E.B., J.D.M., K.M., C.Z., H.S., N.K.A., E.J., M.E., M.I.S., J.F.M.; writing and editing manuscript, C.W., D.C.I.G.; reviewing and editing manuscript, S-J.F., B.K., P.M., E.B., J.D.M., K.M., C.Z., H.S., N.K.A., E.J., M.E., M.I.S., C.C.M., S.M.W., C.C., F.C.H., J.F.M., A.L.H.; funding acquisition, A.L.H., C.W., D.C.I.G.

## SUPPLEMENTARY FIGURE LEGENDS

**Supplementary Figure S1.**
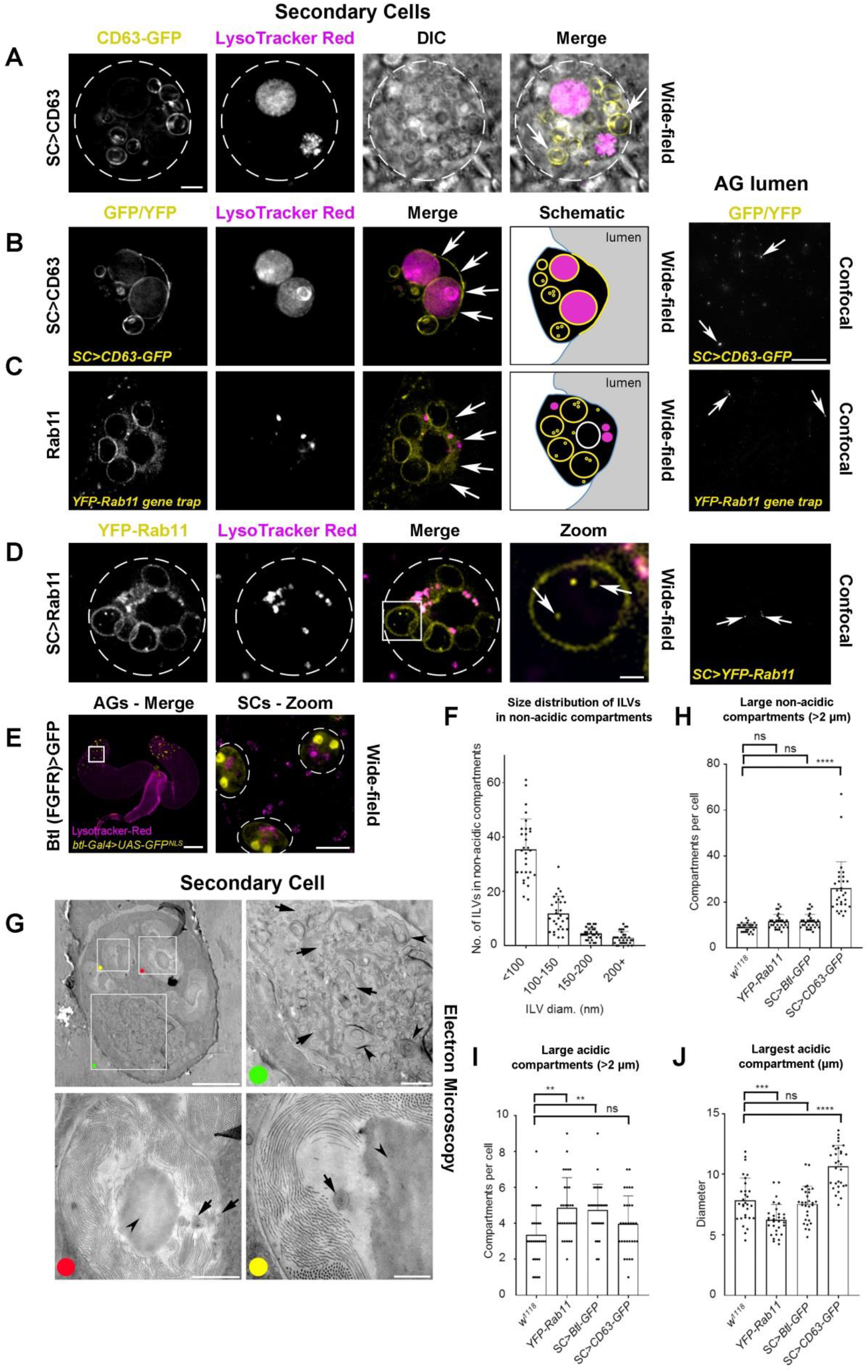
Exosomes Are Made Inside Rab11-Compartments in *Drosophila* Secondary Cells (Related to Figure 1) Panels A-E show wide-field fluorescence images of living fly secondary cells (SCs) and paired accessory glands (AGs; E), and confocal images of fixed AG lumens (B-D). Schematics in Figure 1A have further details of focal planes employed. Acidic compartments are marked by the vital dye LysoTracker^®^ Red (magenta); GFP- and YFP-tagged constructs are shown in yellow. (A) Basal views of an SC, with outline approximated by dashed white circle, including differential interference contrast (DIC) in the two right-hand panels. SC expresses GFP-tagged CD63 (CD63-GFP), which is internalised on intraluminal vesicles (ILVs; see also Figure 1B) in large dense-core granule compartments. The latter have a ‘*fried egg*’ appearance in DIC (arrows) and are Rab11-positive (Corrigan et al., 2014; Redhai et al., 2016). (B) Transverse view of an SC from a fly in which CD63-GFP is expressed specifically in SCs under GAL4/UAS control. CD63-GFP is present on the limiting membrane and ILVs of acidic and non-acidic compartments, in addition to the apical plasma membrane (arrows in merge). Image is also shown schematically. CD63-GFP puncta (arrows) are present in the AG lumen of these flies. (C) Transverse view of an SC expressing a *YFP-Rab11* gene trap, which marks non-acidic compartments, and also the cytosol, but is not trafficked to the apical plasma membrane, (arrows in merge). Image is also shown schematically. Absence of YFP-Rab11 throughout the apical plasma membrane is shown in the Z-stack in Supplementary Movie S2. YFP-Rab11 puncta (arrows) are present at low levels in the AG lumen. (D) Basal view of an SC, with its outline approximated by dashed white circle, from a fly expressing SC-specific YFP-Rab11 (yellow) under GAL4/UAS control. Boxed non-acidic compartment is magnified in Zoom; arrows highlight YFP-Rab11-positive ILVs (Merge) and puncta present at low levels (AG lumen). (E) AGs from a fly expressing nuclear GFP under the control of a well-characterised *btl-GAL4* enhancer trap. Boxed region of AG epithelium containing SCs is enlarged in Zoom, with SC outlines approximated by dashed white circles. Nuclear GFP localisation is specific to binucleate SCs, suggesting *btl* is normally expressed in SCs. (F) Bar chart showing the number of ILVs within specific size ranges in the non-acidic compartments of SCs expressing CD63-GFP. The size of all ILVs was measured in three compartments for three SCs per gland from three independent animals. (G) Transmission electron micrograph of a *w^1118^* non-transgenic fly SC in AG epithelium with boxes showing location of enlarged images, marked by coloured dots. Large compartments lacking dense-core granules (green dot) are equivalent to the acidic compartments seen in live fluorescence imaging (Corrigan et al., 2014). Arrows mark exosome-sized (50-150 nm diameter) vesicles and arrowheads mark complex membranous structures, characteristic of lysosomal compartments. Large non-acidic, dense-core granule compartments (red and yellow dots) have previously been reported to be Rab11-positive (Redhai et al., 2016). Arrows mark ILVs. Arrowhead in each image marks dense core granule, which is surrounded by filamentous material (eg Corrigan et al., 2014). (H) Bar chart shows average number of large (diameter greater than two micrometres), non-acidic dense-core granule compartments, identified using DIC microscopy, in SCs from *w^1118^*, *YFP-Rab11* gene trap, *SC>Btl-GFP* and SC>*CD63-GFP* male flies. Note that the number of these compartments is increased by CD63-GFP expression, as previously reported (Redhai et al., 2016), but not by the other transgenes. (I) Bar chart shows average number of large (diameter greater than two micrometres), LysoTracker Red^®^-positive, acidic compartments (LELs) in different genetic backgrounds. The number of acidic compartments is slightly increased by the *YFP-Rab11* gene trap and *SC>Btl-GFP*. (J) Bar chart shows average size of largest acidic LEL compartment in these genetic backgrounds. The *YFP-Rab11* gene trap slightly reduces largest compartment size, while CD63-GFP increases it. All images and data are from six-day-old males shifted to 29°C at eclosion. Genotypes of flies carrying multiple transgenes are: *w; P[w^+^, UAS-CD63-GFP] P[w^+^, tub-GAL80^ts^]/+*; *dsx-GAL4/+* (A, B, F, H-J), *w; P[w^+^, tub-GAL80^ts^]/+*; *dsx-GAL4/P[w^+^, UAS-YFP-Rab11]* (D), *w; P[w^+^, UAS-GFP^nls^]/+; btl^NP6593^/+* (E), *w; P[w^+^, tub-GAL80^ts^]/+*; *dsx-GAL4/P[w^+^, UAS-btl-GFP]* (H-J). Scale bars in A (5 µm) applies to A-D, and in AG lumen in B (20 µm) applies to B-D. Other scale bars are: 2 µm (D, Zoom), 250 µm (E), 20 µm (E, Zoom), 5 µm (G), 1 µm (G, green and red dots) and 0.5 µm (G, yellow dot). For H-J, values for transgenic flies are compared to *w^1118^*, using one-way ANOVA (n = 10 glands). ***P < 0.001; **P < 0.01; *P < 0.05; n.s. = not significant.

**Supplementary Figure S2.**
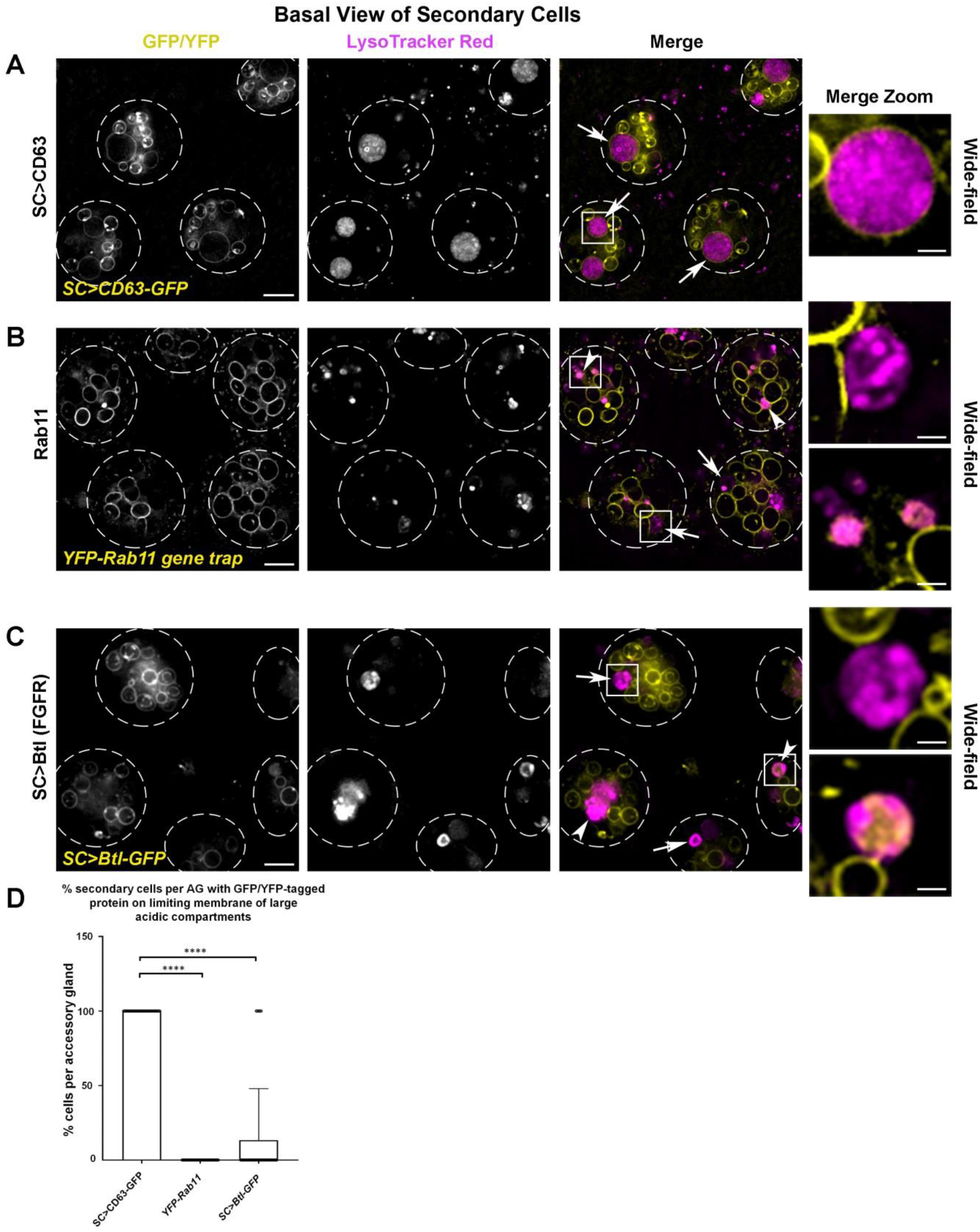
YFP-Rab11 and Btl-GFP Do Not Traffic to the Limiting Membrane of Acidic Compartments in Secondary Cells (Related to Figures 1 and 2) Panels A-C show basal wide-field fluorescence views through living secondary cells (SCs) in same plane as schematic in right panel of Figure 1A, and expressing GFP- and YFP-tagged genes (yellow), with acidic compartments marked by LysoTracker Red^®^ (magenta). (A) CD63-GFP is apparent on the limiting membrane of all large acidic LEL compartments (arrows), when overexpressed in SCs. Boxed region in Merge is magnified in Merge Zoom (see also quantification in D). (B) YFP-Rab11 is never found on the limiting membrane of acidic compartments in gene trap males (see also D). Boxed regions are magnified in Zoom. Some compartments lack internal YFP-Rab11 (arrows; upper Zoom panel), while others have fluorescent luminal content (arrowheads; lower Zoom panel). (C) Btl-GFP is mostly absent from the limiting membrane of acidic compartments, when overexpressed in SCs (arrows). Boxed regions are magnified in Zoom. Some compartments lack internal Btl-GFP (arrows; upper Zoom panel), while others have fluorescent luminal content (arrowheads; lower Zoom panel). (D) Bar chart showing proportion of SCs containing a large acidic LEL compartment with CD63-GFP, YFP-Rab11 and Btl-GFP on its limiting membrane. All images are from six-day-old male flies shifted to 29°C at eclosion. Genotypes of flies carrying multiple transgenes are: *w; P[w^+^, UAS-CD63-GFP] P[w^+^, tub-GAL80^ts^]/+*; *dsx-GAL4/+* (A): *w; P[w^+^, tub-GAL80^ts^]/+*; *dsx-GAL4/P[w^+^, UAS-btl-GFP]* (C). Values in D compared using the Kruskal-Wallis test (n = 10 AGs). ****P < 0.0001. Scale bar in A-C (10 µm) and and in Zoom (2 µm).

**Supplementary Figure S3.**
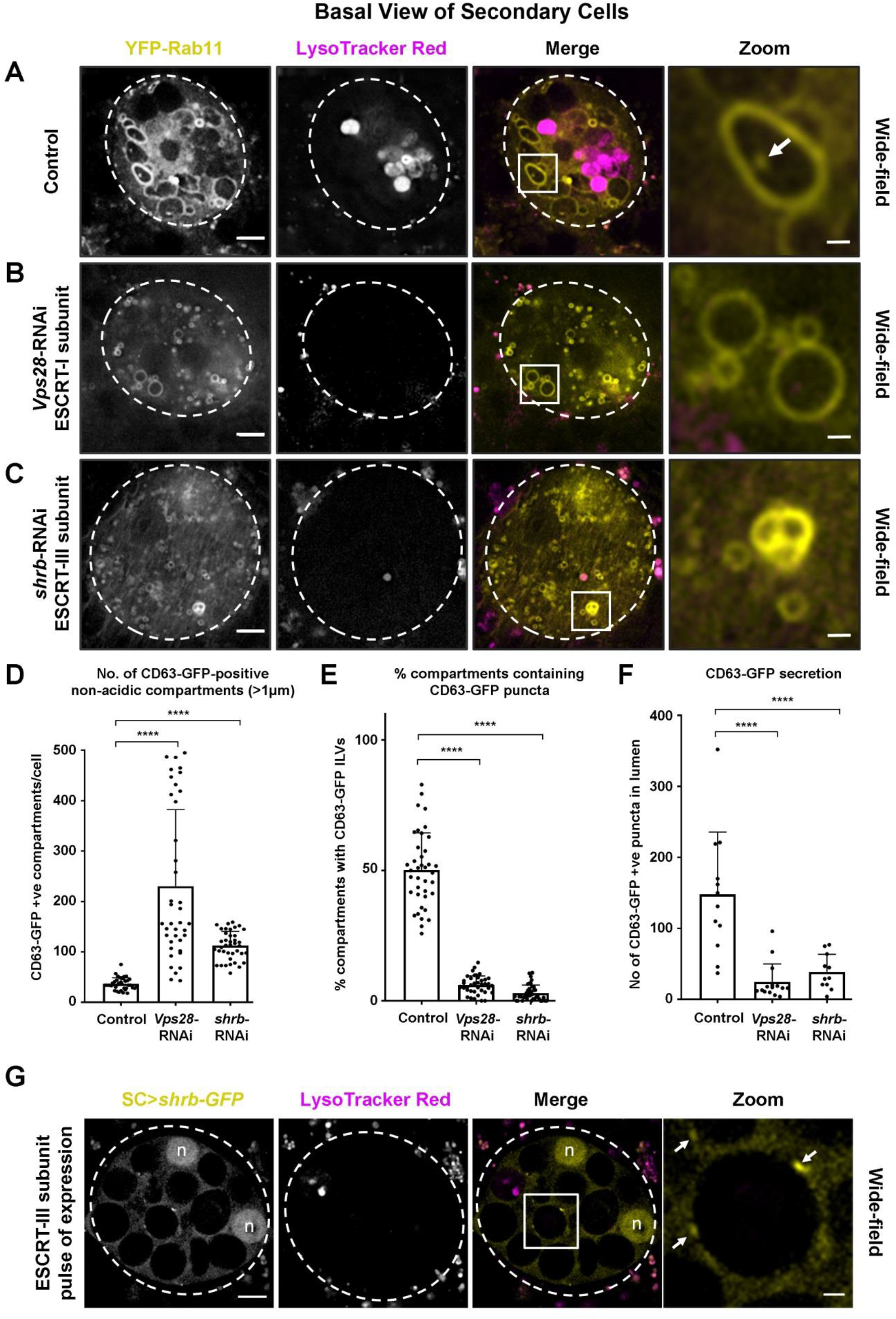
ESCRTs Regulate Exosome Biogenesis in the Rab11-Positive Compartments of *Drosophila* Secondary Cells (Related to Figure 2) Panels A-C show basal wide-field fluorescence views through living secondary cells (SCs) expressing the *YFP-Rab11* gene trap. Acidic compartments are marked with LysoTracker Red^®^ (magenta). Boxed non-acidic compartment in Merge is magnified in Zoom. (A) SC with no RNAi expressed (control). Arrow marks YFP-Rab11-positive internal ILV puncta (Zoom). (B) SC also expressing RNAi targeting ESCRT-I component, *Vps28*. Number of Rab11-positive compartments is increased, but few contain ILVs (Zoom). (C) SC also expressing RNAi targeting ESCRT-III component *shrb*. Number of Rab11-positive compartments is increased, but few contain ILVs (Zoom). (D) Bar chart showing the number of large (greater than one micrometre in diameter) non-acidic compartments marked by CD63-GFP in control and *ESCRT* knockdown SCs. Data from 39 SCs (three per gland) are shown. (E) Bar chart showing the proportion of large (greater than one micrometre in diameter) non-acidic compartments containing CD63-GFP-positive ILVs in control and *ESCRT* knockdown SCs. Data from 39 SCs (three per gland) are shown. (F) Bar chart showing the total number of CD63-GFP fluorescent puncta in three Z-planes from the lumen of AGs following *ESCRT* knockdown in SCs, compared to control SCs (n = 10 AG lumens). (G) SC, after 4 h pulse of Shrb-GFP (yellow). Arrows highlight Shrb-GFP localisation on the limiting membranes of large non-acidic compartments (Zoom). Complete view of Z-stack is shown in Movie S3. Nuclear staining (marked ‘n’) of these binucleate cells is non-specific. All images are from six-day-old male flies shifted to 29°C at eclosion, except for G, where flies were cultured at 25°C for six days before 29°C pulse. Genotype of male in G is *w; P[w+, UAS-shrb-GFP]/P[w^+^, tub-GAL80^ts^]; dsx-GAL4/+*. Scale bar in A-C, G (5 µm) and in A-C, G Zoom (1 µm). Data were analysed by one-way ANOVA. ****P < 0.0001.

**Supplementary Figure S4.**
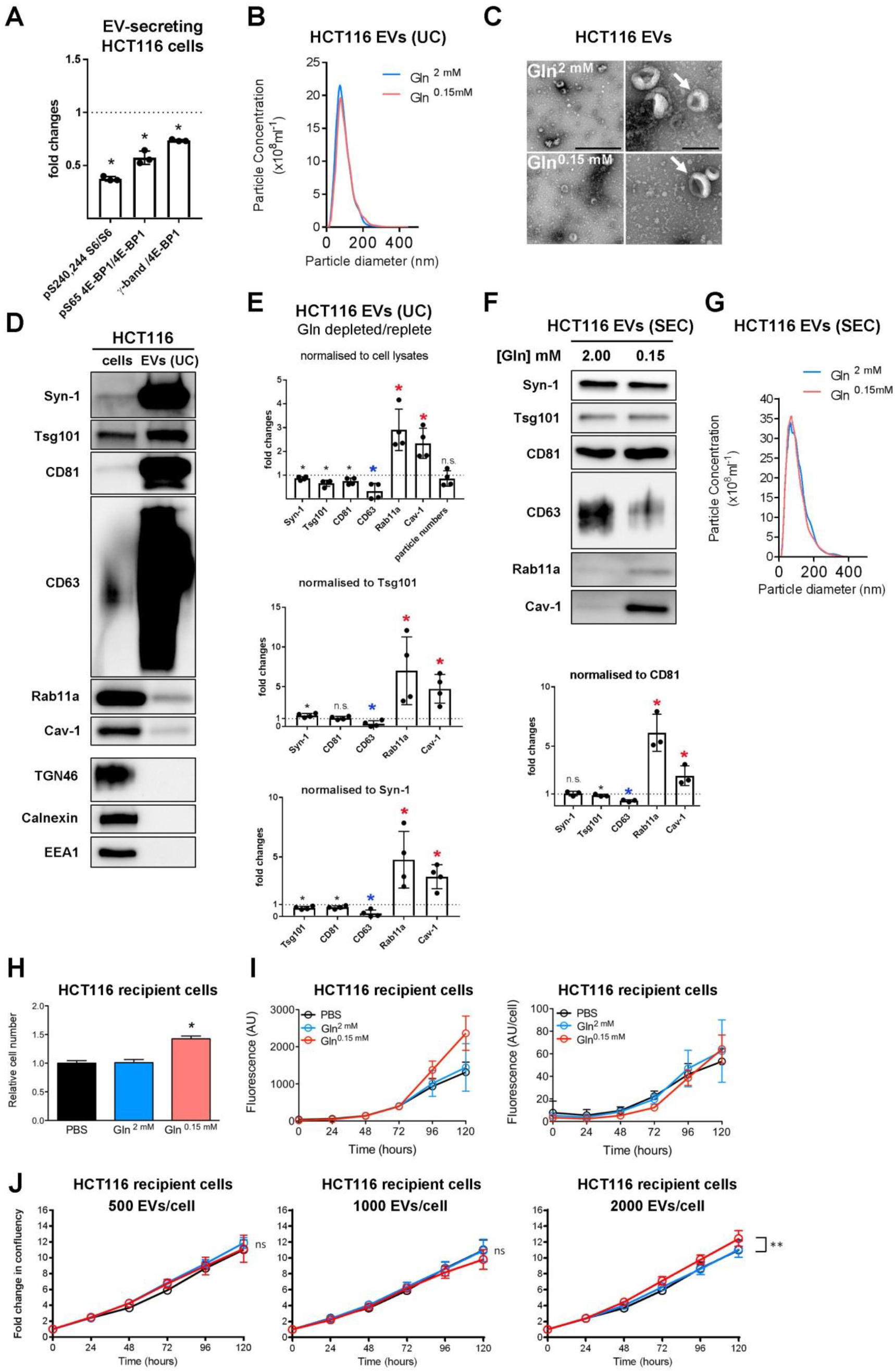
Effects of Glutamine Depletion on HCT116 Cells and Their EVs (Related to Figures 4 and 5) Panels show the effect of different treatments on HCT116 colorectal cancer cells and on the EVs they secrete. (A) Bar chart showing relative levels of phosphorylation of mTORC1 downstream readouts, S6 and 4E-BP1 (measured as a ratio of p-S65-4E-BP1 to pan 4E-BP1 or of the γ-phosphorylated form of 4E-BP1 relative to total 4E-BP1), following growth of cells for 24 h in glutamine-depleted (2.00 mM) versus -replete (0.15 mM) conditions. Data are from triplicate experiments, analysed as in Figure 4A. (B) Nanosight Tracking Analysis of EV size and number for diluted samples (normalised to cell lysate protein levels) produced from ultracentrifugation (UC) of medium from cells cultured in glutamine-replete and -depleted conditions for 24 h, as in Figure 4B. (C) Electron micrographs (EMs) of EV samples from glutamine-replete and -depleted cells as analysed in (B). Arrows indicate representative EVs with the characteristic cup-shaped morphology previously reported for EM samples. Scale bars represent 500 nm (left panels) and 200 nm (right panels). (D) Comparative western analysis of putative exosome and non-exosome markers in HCT116 cell lysate (cells) and EV preparation produced by UC from cells grown in glutamine-replete conditions (EVs). Lanes are loaded with equal amounts of protein. Note that unlike other exosome markers, Rab11a and Cav-1 are present, but not enriched in EVs, while Golgi (TGN46), ER (Calnexin) and early endosome (EEA) markers are not found in EVs. (E) Bar charts show changes in levels of putative exosome proteins in EVs produced by UC from cells grown in glutamine-depleted versus -replete conditions and analysed as in Figure 4B. Protein quantities in each sample were normalised to cell lysate protein levels (top), Tsg101 and Syn-1 (bottom) in the three graphs. (F) Western blot analysis of EV preparations isolated from HCT116 cells cultured in glutamine-replete and -depleted conditions for 24 h using size-exclusion chromatography (SEC; fractions two to five). EV loading was normalised to protein levels in cell lysates. Bar chart shows relative levels of putative exosome markers normalised to CD81. (G) Nanosight Tracking Analysis of EV size and number for samples produced as in (F). (H) Number of DAPI-stained cells after 120 h culture in serum-depleted conditions (1% serum), following pre-incubation either with EVs from HCT116 cells cultured in glutamine-replete or -depleted conditions for 24 h, or with vehicle (PBS). Bar charts derived from three independent experiments. (I) Overall levels of apoptosis, measured in arbitrary units (AU), of fluorescence, and levels of apoptosis per cell (right hand graph) detected in HCT116 cells by analysis of Caspase-3 and −7 activities, following treatments with EVs isolated as in (H) or with vehicle (PBS). (J) Growth curves for HCT116 recipient cells in 1% serum conditions following 30 min pre-incubation with 0.5 x 10^3^, 1.0 x 10^3^ and 2.0 x 10^3^ EVs per cell, isolated by SEC from glutamine-replete and -depleted HCT116 cells or PBS. Bar charts derived from three independent experiments; *P < 0.05; **P < 0.01; n.s. = not significant.

**Supplementary Figure S5.**
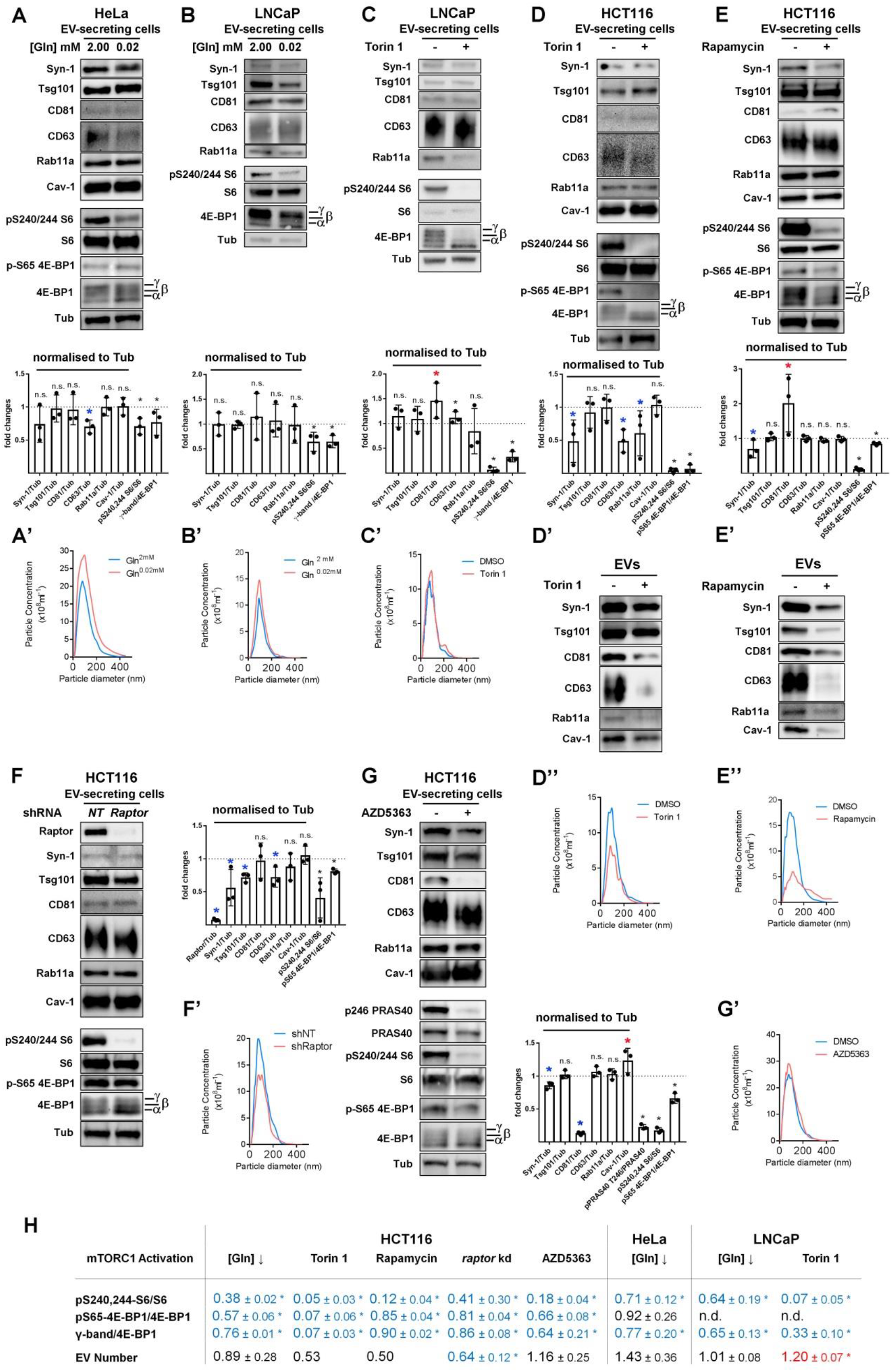
Changes in EV-Secreting Cell Proteins and EVs Following Glutamine Depletion and Reduction in PI3K/Akt/mTORC1 Signalling in Three Human Cancer Cell Lines (Related to Figure 4) Panels show western blot analyses of cell and EV proteins, as well as Nanosight Tracking Analysis of EV size and number. Bar charts indicate abundance of putative exosome proteins relative to Tubulin in cell lysates and relative to CD81 in EVs. (A) With relevance to EVs shown in Figure 4C, western blot analysis of lysates from HeLa cells cultured in glutamine-replete (2.00 mM) and -depleted (0.02 mM) media for 24 h. Total protein levels were reduced by 19 ± 4% after glutamine depletion. (A’) Nanosight Tracking Analysis for EV samples produced as in Figure 4C. (B) With relevance to EVs shown in Figure 4D, western blot analysis of lysates from LNCaP cells cultured in glutamine-replete (2.00 mM) and -depleted (0.02 mM) media for 24 h. Total protein levels were unaffected (100 ± 11% of control values) after glutamine depletion. (B’) Nanosight Tracking Analysis for EV samples produced as in Figure 4D. (C) With relevance to EVs shown in Figure 4E, western blot analysis of lysates from LNCaP cells cultured in the presence or absence of 120 nM Torin 1 for 24 h. Total protein levels were reduced by 19 ± 1% following drug treatment. (C’) Nanosight Tracking Analysis for EV samples produced as in Figure 4E. (D) Western blot analysis of lysates from HCT116 cells cultured in the presence or absence of 100 nM Torin 1 for 24 h. Total protein levels were reduced by approximately 20% following drug treatment. (D’) Western blot analysis of EV preparations isolated by size-exclusion chromatography from HCT116 cells cultured in the presence or absence of 100 nM Torin 1 for 24 h. EV loading was normalised to protein level in cell lysates. (D”) Nanosight Tracking Analysis for EV samples produced as in (D’). (E) Western blot analysis of lysates from HCT116 cells cultured in the presence or absence of 10 nM rapamycin for 24 h. Total protein levels were reduced by 7 ± 2% following drug treatment. (E’) Western blot analysis of EV preparations from HCT116 cells isolated by ultracentrifugation from HCT116 cells cultured in the presence or absence of 10 nM rapamycin for 24 h. EV loading was normalised to protein level in cell lysates. (E”) Nanosight Tracking Analysis for EV samples produced as in (E’). (F) With relevance to EVs shown in Figure 4F, western blot analysis of lysates from HCT116 cells subjected to four days of *raptor* or non-targeting (*NT*) shRNA knockdown. Total protein levels were not significantly altered by knockdown versus control. (F’) Nanosight Tracking Analysis for EV samples produced as in Figure 4F. (G) With relevance to EVs shown in Figure 4G, western blot analysis of lysates from HCT116 cells cultured in the presence or absence of 3 µM AZD5363 for 24 h. Total protein levels were reduced by 12 ± 5% following drug treatment. Note that phosphorylation of PRAS40, an Akt target, is reduced by drug treatment. (G’) Nanosight Tracking Analysis for EV samples produced as in Figure 4G. (H) Table summarising relative EV secretion, and relative activity of mTORC1 assessed by analysing levels of 4E-BP1 and S6 phosphorylation under conditions shown in panels A to G, and in Figure 4A: significantly decreased levels versus control are marked in blue and increased levels in red. Note strong inhibition of S6 phosphorylation in Torin 1- and rapamycin-treated HCT116 cells, which is associated with low levels of EV and exosome secretion. Bar charts derived from three independent experiments: **P < 0.01; *P < 0.05; n.s. = not significant.

**Supplementary Figure S6.**
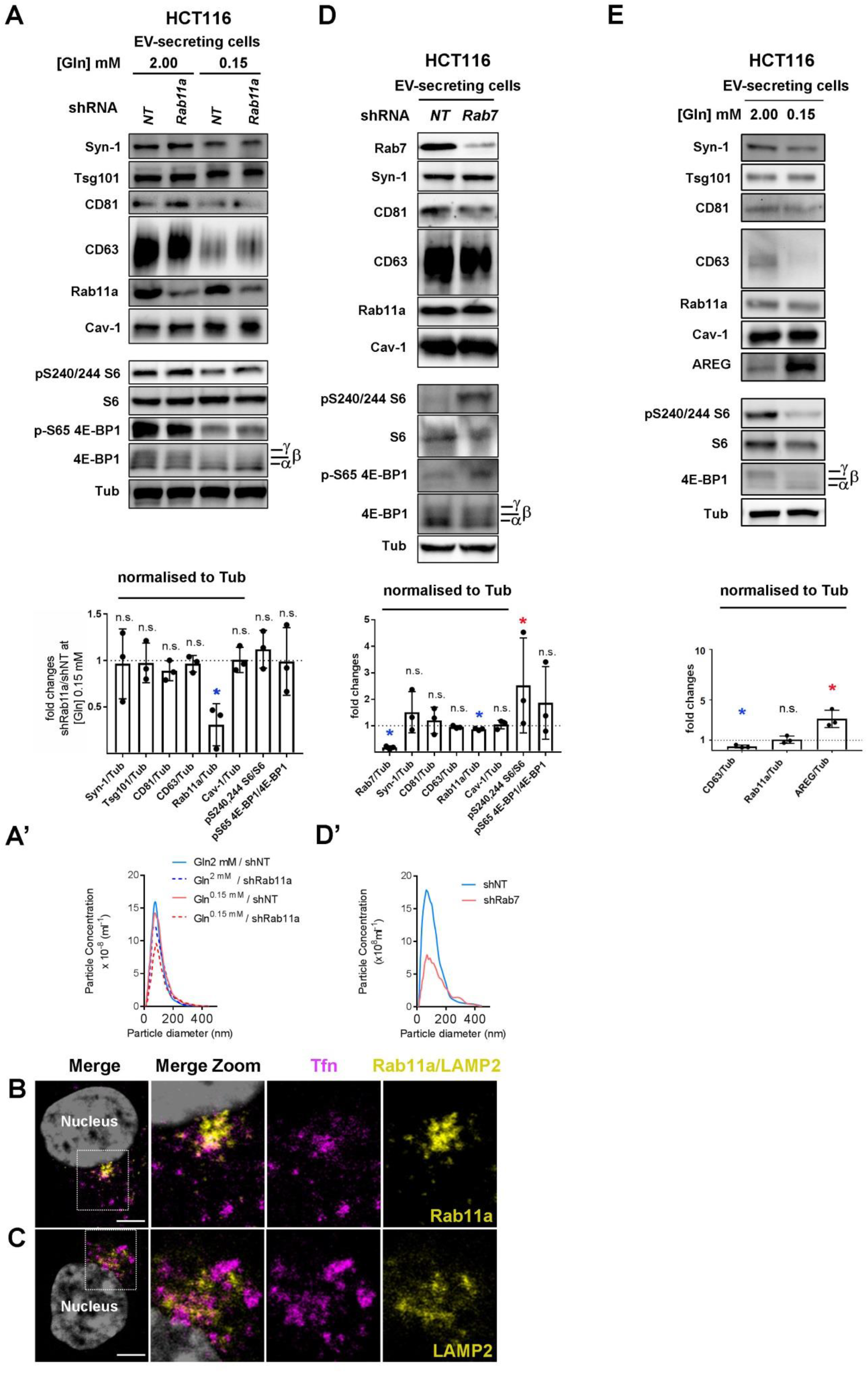
Analysis of EVs and Secreting Cells from Control and Knockdown HCT116 Colorectal Cancer Cells, and Effects of ERK Inhibitor and Anti-AREG Antibody on HCT116 Recipient Cell Growth (Related to Figures 5 and 6) Panels A, D and E show western blot analysis of cell lysate proteins, and Nanosight Tracking Analysis of EV size and number in A’ and D’. Bar charts indicate putative exosome protein levels normalised to Tubulin on western blots. (A) With relevance to EVs shown in Figure 5C, western analysis of lysates from EV-secreting cells with and without knockdown of *Rab11a* in glutamine-replete and -depleted conditions. Total protein levels were increased by 1 ± 3% following knockdown. (A’) NanoSight Tracking Analysis of EVs isolated from HCT116 cells with and without knockdown of *Rab11a*. (B) With relevance to the protein uptake assays shown in Figure 5D, HCT116 cells stained with an anti-Rab11a antibody (yellow), following uptake of fluorescent Alexa-488-conjugated Tfn (magenta); boxed region enlarged in Zoom. Some Tfn co-localises with Rab11a. DAPI (grey) stains nucleus. (C) Cells stained with an anti-LAMP2 antibody (yellow), following uptake of fluorescent Alexa-488-conjugated Tfn (magenta); boxed region enlarged in Zoom. There is little overlap between Tfn and LAMP2. DAPI (grey) stains nucleus. (D) With relevance to EVs shown in Figure 5F, western analysis of proteins isolated from EV-secreting HCT116 cells with and without knockdown of *Rab7*. (D’) NanoSight Tracking Analysis of EVs isolated from HCT116 cells with and without knockdown of *Rab7*. (E) With relevance to EVs shown in Figure 6C western blot analysis of cell lysates from HCT116 cells cultured in glutamine-replete (2.00 mM) and glutamine-depleted (0.15 mM) medium for 24 h. Gel was loaded with equal protein amounts. The activity of mTORC1 was assessed via phosphorylation of S6 and 4E-BP1. Bar chart shows the abundance of selected exosome/EV proteins relative to tubulin in these lysates. Bar charts derived from three independent experiments: *P < 0.05; n.s. = non-significant. Scale bars in B, C (5 µm).

**Supplementary Figure S7.**
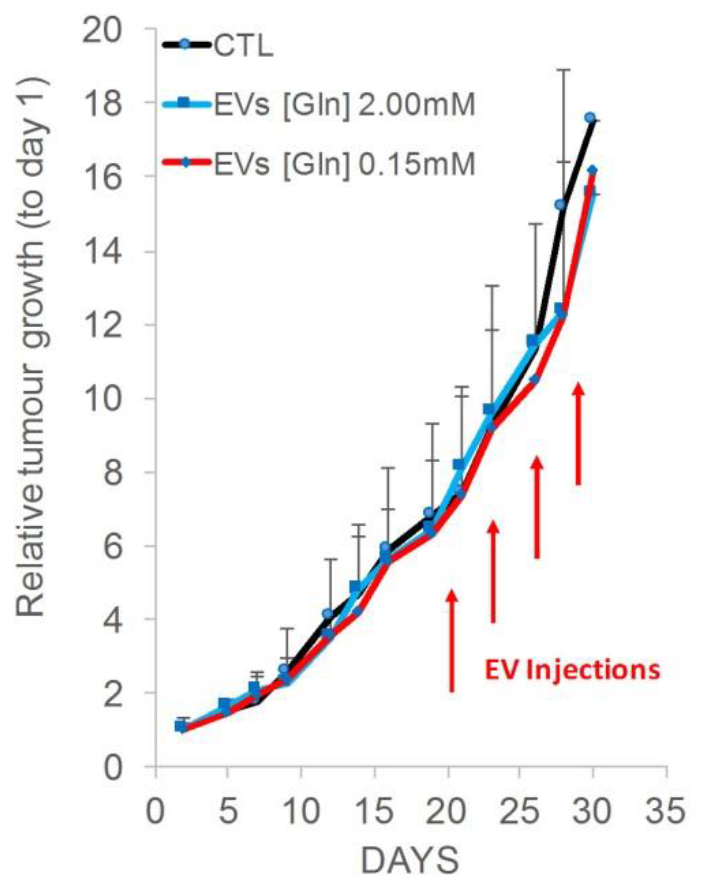
Overall Growth of HCT116 Tumours Grown in Xenograft Mouse Model is Not Affected by EV Injection Over Nine Days (Related to Figure 7). The size of HCT116 tumours grown in xenograft mouse models, generated and treated as in Figure 7, was measured every two to three days. Arrows mark times of the four EV injections in to the tumour. There was no significant difference in size for any of the treatments: PBS injection control, EVs isolated from glutamine-replete control or glutamine-depleted HCT116 cells.

## SUPPLEMENTARY MOVIES

**Supplementary Movie S1. Intraluminal Vesicles Are Present in Non-Acidic Compartments of *Drosophila* Secondary Cells (Related to Figure 1)**

Movie of Z-stack generated from super-resolution 3D-SIM images of a living SC expressing CD63-GFP (yellow) and labelled with LysoTracker Red^®^ (magenta). It focuses on a single non-acidic compartment, which has intraluminal vesicles (ILVs) of varying size arranged in at least four clusters. ILV size is quantified in Figure S1F.

Genotype of fly is *w; P[w^+^, UAS-CD63-GFP] P[w^+^, tub-GAL80^ts^]/+*; *dsx-GAL4/+*.

Scale bar is 1 µm. The Z-stack includes 51 sections at 0.125 µm intervals.

**Supplementary Movie S2. Rab11 is Not Trafficked to the Apical Plasma Membrane of *Drosophila* Secondary Cells (Related to** Figure 1)

Movie of Z-stack generated from wide-field fluorescence transverse images of a living SC of a male fly expressing a YFP-Rab11 gene trap (yellow). Figure S1C shows one of these images. Acidic compartments are marked by LysoTracker Red^®^ (magenta). Unlike CD63-GFP, YFP-Rab11 is not localised to the apical plasma membrane. Punctate intraluminal vesicles are marked by YFP-Rab11 inside large Rab11-compartments (more clearly seen in Figure S1C). YFP-Rab11 is also observed in the cytosol.

Scale bar is 5 µm. The Z-stack includes 27 sections at 0.5 µm intervals.

**Supplementary Movie S3. Shrb-GFP Accumulates in Microdomains at the Surface of Large Non-Acidic Compartments in *Drosophila* Secondary Cells (Related to** Figure S3G)

Movie of Z-stack generated from wide-field fluorescence images of a living SC after a 4 h pulse of Shrb-GFP. Acidic compartments are marked by LysoTracker Red^®^ (magenta). Note the accumulation of Shrb-GFP in microdomains at the surface of non-acidic and acidic compartments. Figure S3G shows a single Z-plane from this movie.

Scale bar is 5 µm. The Z-stack includes 56 sections at 0.2 µm intervals.

